# Patterns of change in regulatory modules of chemical reaction systems induced by network modification

**DOI:** 10.1101/2023.11.24.567370

**Authors:** Atsuki Hishida, Takashi Okada, Atsushi Mochizuki

## Abstract

Cellular functions are realized through the dynamics of chemical reaction networks formed by thousands of chemical reactions. Numerical studies have empirically demonstrated that small differences in network structures among species or tissues can cause substantial changes in dynamics. However, the existence of a general principle for behavior changes in response to network structure modifications is not known. The chemical reaction system possesses substructures called buffering structures, which are characterized by a certain topological index being zero. It was proven that the steady-state response to modulation of reaction parameters inside a buffering structure is localized in the buffering structure [1, 2, 3]. In this study, we developed a method to systematically identify the loss or creation of buffering structures induced by the addition of a single degradation reaction from network structure alone. This makes it possible to predict the qualitative and macroscopic changes in regulation that will be caused by the network modification. This method was applied to two reaction systems: the central metabolic system and the mitogen-activated protein kinases (MAPK) signal transduction system. Our method enables identification of reactions that are important for biological functions in living systems.

**Significance Statement:** Dynamics of complex biological systems have been understood using mathematical models[4, 5, 6, 7]; however, various assumptions are necessary to construct models for large networks such as those found in the life sciences, and discussing the relation between the network structure and its behavior is difficult. Our study using structural sensitivity analysis [1, 2, 3] shows that directly discussing changes in the structure and behavior of a reaction network without making assumptions about kinetics is possible. This enabled the prediction of changes in behavior depending on whether a reaction is present or not and identify reactions that are essentially important for biological behavior.

## Introduction

### Differences in network structure induce differences in cellular functions

Within cells, a large number of enzymes catalyze their specific reactions simultaneously, and these reactions form complex reaction networks, such as metabolic systems or signal transduction pathways. Cellular functions arise from the dynamics of such complex networks, and cells regulate network dynamics by modulating the activity or expression level of enzymes [8, 9]. In the history of biosciences, structures of reaction networks responsible for various cellular functions have been experimentally identified. For example, cells obtain chemical energy via a series of reactions in the tricarboxylic acid cycle (TCA) or regulate gene expression via signal transduction networks such as the MAPK pathway when they receive extracellular signals [10, 11, 12, 13]. Revealing the mechanism for controlling the dynamics of those networks will lead to understanding how cells adapt to their environments by altering cellular functions.

During the course of evolution, species have acquired different networks that serve distinct cellular functions. Cancer cells are also known to have reaction networks with a different structure from normal cells [8, 14, 15]. Those structural differences may alter cellular functions derived from network dynamics, and such alterations can be advantageous for the survival of organisms or cancer cells [14, 15, 8]. However, it is not clear how these structural differences influence the dynamics of reaction systems and biological functions of organisms [16]. Databases of biological networks provide information regarding the network structures of different cell types; however, they do not provide insight into the differences in dynamics of these networks [17, 18, 19]. Therefore, to date, no general rules have been obtained regarding how structural differences in reaction networks affect cellular functions.

### Structural sensitivity analysis: a mathematical method to determine the sensitivity of the system

Structural sensitivity analysis (SSA), which we previously developed, is a mathematical method for determining the response of a steady state to a change in enzyme activity based on the structure of a reaction network [1, 3] (see the Methods section B. for the summary of the method). In general, the dynamics of reaction systems are quantitatively determined only when we have full knowledge of reactions, but experimentally determining the rate functions of those reactions is difficult when the system is large and complicated. SSA is a model-free method by which the steady-state response to changes in reaction parameters is determined without assuming reaction functions or parameter values.

The concept of a “buffering structure,” derived from SSA, is crucial in understanding the responses of a system to a parameter change based on the network structure. A substructure in a reaction network, which consists of a subset of chemicals and reactions, is called a buffering structure when it satisfies the following two conditions: (i) The substructure contains all reactions whose rate depends on the concentration of chemicals within it. (ii) The index defined from elements in the substructure (later defined in Eq. 1) The first condition is referred to as the “output-completeness”. See [2, 3] or the Methods section C. for the precise definitions for the index. It is mathematically proven that the steady-state response to the activation or inhibition of reactions within a buffering structure is confined to the substructure. From a biological perspective, a buffering structure is understood as the origin of modularity in the regulation of cellular functions generated from reaction systems because ranges of influences of modulation to the activity or expression level of enzymes are restricted in buffering structures. As a result, buffering structures can be independently regulated from other parts in the network through the activation or inhibition of reactions inside them.

For example, in the network shown in Fig. 1A, the substructure ({*D* }, {6}) is a buffering structure because it satisfies output-completeness, and its index is zero. This means that the activation of reaction

**Figure 1:**
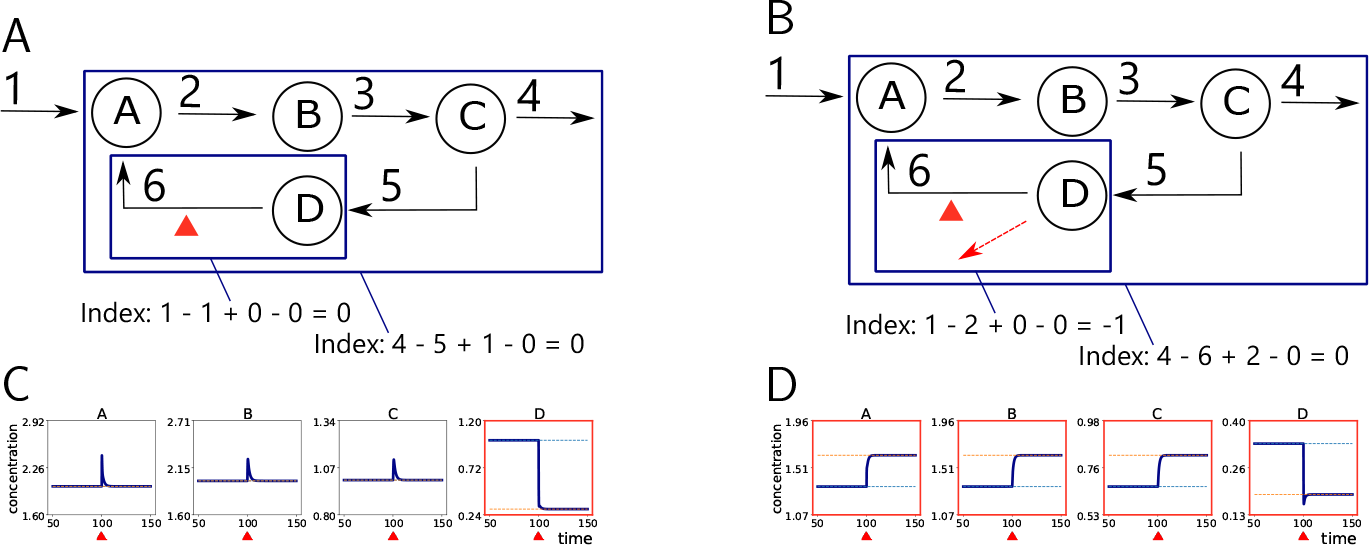
(A) Reaction network constructed from four chemicals A, B, C, and D, and six reactions. Solid lines indicate chemical reactions. Blue rectangles indicate buffering structures. The reaction indicated by the red arrowhead is activated in the numerical simulation shown in (C). (B) A reaction network with the outflow of D added to (A). (C) The result of numerical simulation in the network (A). In this simulation, we assumed mass-action kinetics; *r*_1_ = *k*_1_, *r*_2_ = *k*_2_*x*_*A*_, *r*_3_ = *k*_3_*x*_*B*_, *r*_4_ = *k*_4_*x*_*C*_, *r*_5_ = *k*_5_*x*_*C*_, *r*_6_ = *k*_6_*x*_*D*_. All reaction parameters are set to 2.5. The dynamics reached a steady state at *t* = 100, and then the parameter of reaction 6 was increased from 2.5 to 8.0. Orange and green dotted lines indicate concentration values at the steady state before and after the perturbation, respectively. The frame of the numerical simulation result is colored red if the steady state value differs before and after the perturbation to the reaction parameter. (D) The result of numerical simulation in the network (B) based on mass-action kinetics. All reaction parameters are set to 2.5. At *t* = 100, the parameter of reaction 6 was increased from 2.5 to 8.0. Orange and green dotted lines indicate concentration values at the steady state before and after the perturbation, respectively. The frame of the numerical simulation result is colored red if the steady state value differs before and after the perturbation to the reaction parameter.

6 affects only the concentration of D at a steady state, which is confirmed by the numerical simulation shown in Fig. 1C. In this simulation, we assumed mass-action kinetics, where each rate function is given by the product of a parameter *k*_*i*_ and substrate concentrations powered by stoichiometric coefficients. In contrast, the new network shown in Fig. 1B has a different sensitivity from that of the original network when a degradation reaction of D is added to the network. The activation of reaction 6 in turn affects the concentration of all chemicals in the network, as shown in Fig. 1D. This is because the index of the output-complete substructure ({*D* }, {6, *outflow*-*of* -*D*}) is -1, and it is not a buffering structure. Although the index of the substructure ({*D*}, {6}) in the new network is 0, it is not a buffering structure because the substructure does not satisfy the output-completeness. In the new network, the smallest buffering structure containing reaction 6 is the substructure ({*A, B, C, D* }, {2, 3, 4, 5, 6, *outflow*-*of* -*D* }). This result illustrates that a structural alteration of a reaction network can cause a qualitative change in the sensitivity of the system, and it can be captured by the loss or creation of buffering structures.

In this study, we investigated how the sensitivity of a reaction network is affected by structural modification. Specifically, we considered the addition of an “outflow” reaction (i.e., a reaction for which substrates exist and products do not explicitly exist in the network) and proved that such addition can lead to the loss of existing buffering structures or the creation of new buffering structures in the system. The effects of modulation to reaction parameters inside the substructure extend to the outside when a substructure that used to be a buffering structure no longer satisfies the conditions for being a buffering structure. The creation of a new buffering structure indicates that the effects of modulation to reaction parameters inside the substructure are confined inside itself in the new network. In other words, such a small difference in network structure can result in qualitative and macroscopic changes in the regulations of the reaction system.

Since buffering structures are output-complete substructures with zero index, by examining the index change for each of all output-complete substructures in the system, it is possible to identify all the loss or creation of buffering structures. This idea allows us to predict the qualitative and macroscopic changes in regulation that will occur in the system when an outflow reaction is added to the reaction network. We utilized this result in reverse to identify structural modifications (i.e., the addition of a new reaction) that bring qualitative and macroscopic changes in the regulation of the system. By applying this method to the central metabolic system and MAPK signal transduction system, we evaluated the effect of specific structural modifications on the energy metabolism and signal transduction. We also identified reactions important for the TCA cycle to be a buffering structure. The rule we obtained has significant implications for understanding the dynamics of living systems.

## Result

### Changes in the index of output-complete substructures caused by the addition of an outflow

We investigated how the sensitivity of a reaction network is affected by the structural modification of the network. A reaction system before a modification is called an “original system” denoted by Γ. *M* (Γ) and *R*(Γ) are the sets of chemicals and reactions in Γ, respectively. Among possible network alterations, in this study, we considered the addition of a reaction whose substrate exists and products do not exist in the original system and assessed the effect of the structural modification on the sensitivity. This reaction can be graphically represented as an edge from a single node with no target nodes; we called it an “outflow”.

The biological interpretation of an outflow may be the degradation of a substrate, transport of a substrate from the system to the outside, or a reaction from a substrate in the original system into another chemical excluded from the system. The presence/absence of outflows may not be explicitly stated in databases and tends to be subjectively modeled by theoretical researchers. Depending on the time scale of interest, outflows may be appropriately interpreted as present or absent. Conversely, the presence/absence of outflows has a large effect on the behavior of a reaction system. Although we will focus on the effects of adding outflows, other types of network alternation will be briefly discussed in the Discussion section. While inflows are also biologically important and necessary, their effects are usually understandable in an intuitive way (e.g., see [1] for sensitivity associated with inflows). We compared the sensitivity of Γ and that of the new system Γ^*′*^ (i.e., the network obtained by adding the outflow from chemical *m*^*^ to Γ).

We analyzed the effect of adding an outflow of *m*^*^ to the original system from the viewpoint of changes in buffering structures in the system. The existence of an outflow may induce the creation of new buffering structures or the loss of existing buffering structures, which should cause a large change in the sensitivity. Such large changes can be captured by focusing on index change of “output-complete” substructures.

For an output-complete substructure *γ*, the index of the substructure is defined as

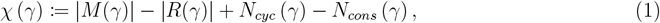

where |*M* (*γ*) |, |*R*(*γ*) |, *N*_*cyc*_ (*γ*), and *N*_*cons*_ (*γ*) represent the numbers of chemicals, reactions, cycles, and conserved quantities, respectively. We define the number of cycles as the number of independent steady-state fluxes flowing inside the substructure. An output-complete substructure *γ* is a buffering structure if *χ*(*γ*) = 0 holds (see [2, 3] or Eq. 4, 5 for the precise definitions of *N*_*cyc*_ (*γ*) and *N*_*cons*_ (*γ*)).

For each of output-complete substructures, we studied the index change by the addition of outflow of *m*^*^. For an output-complete substructure *γ* in the original system Γ constructed by a set of chemicals *M* (*γ*) and a set of reactions *R*(*γ*), the corresponding output-complete substructure in the new system Γ^*′*^, denoted as *γ*^*′*^ with chemicals *M* (*γ*^*′*^) and reactions *R*(*γ*^*′*^), is defined as follows. The set of chemicals *M* (*γ*^*′*^) is identical to *M* (*γ*). If *M* (*γ*) (or *M* (*γ*^*′*^)) does not contain the target chemical *m*^*^, then the reaction set *R*(*γ*^*′*^) is equal to *R*(*γ*). If *M* (*γ*) contains *m*^*^, then *R*(*γ*^*′*^) includes the new outflow of *m*^*^ in addition to *R*(*γ*). Because *γ* satisfies output-completeness, *γ*^*′*^ also does.

We identified changes in the sensitivity of the system by comparing the values of *χ* (*γ*) and *χ* (*γ*^*′*^), particularly focusing on whether either of them takes the zero value. If any output-complete substructures satisfy *χ* (*γ*) − *χ* (*γ*^*′*^) ≠ 0, the addition of the outflow can qualitatively change the sensitivity.

We proved that the change in the index of each output-complete substructure in an original system Γ is categorized into five cases depending on three conditions: (i) whether the substructure includes *m*^*^, (ii) whether it contains a conserved quantity composed of *m*^*^, and (iii) whether a new cycle is generated in it by the addition of the outflow (Fig. 2A, B). See the Methods section D. for the proof. An example of each case is shown in Fig. 2C.

**Figure 2:**
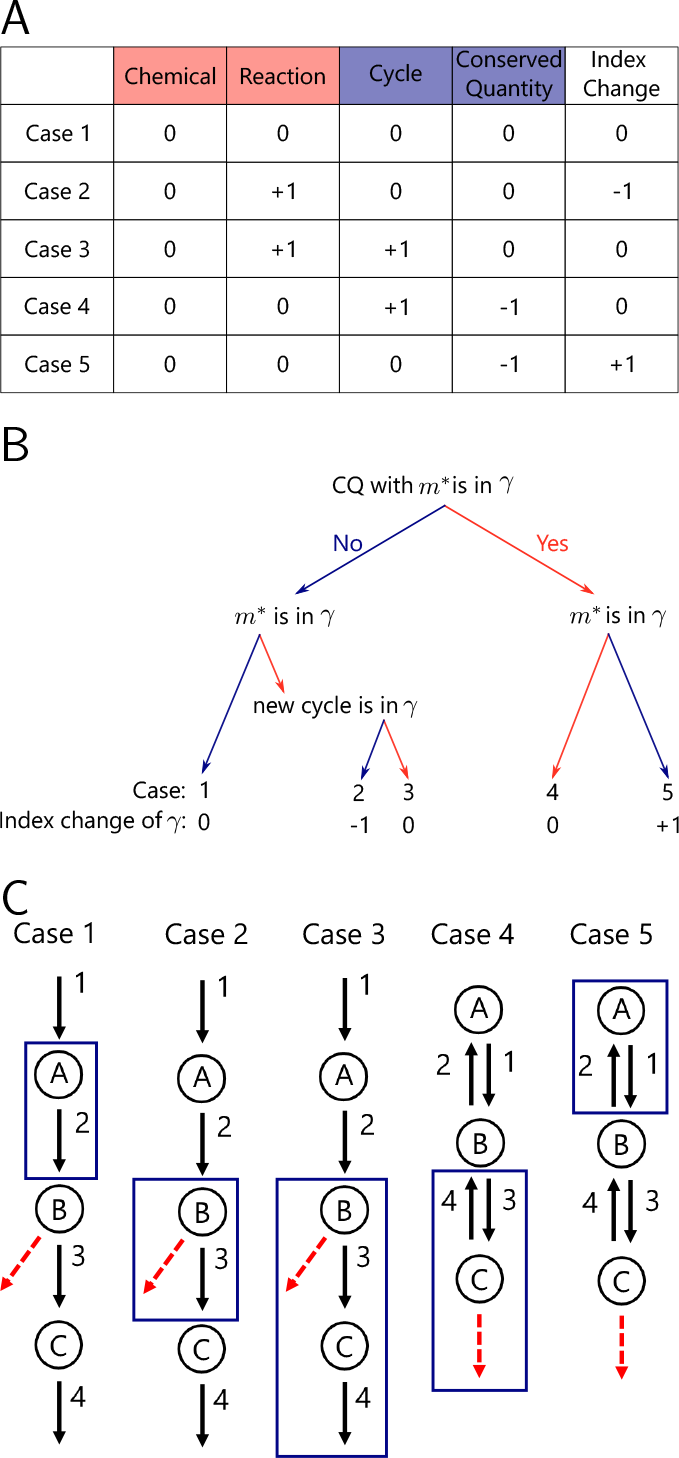
(A) Change in the index of an outputcomplete substructure *γ* under the addition of an outflow is classified into five patterns, and the amount of change is limited only to − 1, 0, and +1. Which pattern the index change of *γ* follows can be determined by three conditions: whether *γ* contains the substrate of the outflow, whether the outflow creates a new cycle with reactions in *R*(*γ*), and whether *M* (*γ*) contains a chemical that constructs a conserved quantity of Γ with *m*^*^. (B) Example network and substructure of each case. The blue rectangle indicates a substructure *γ*, and the red arrow indicates an outflow of *m*^*^ added to Γ. In cases 2, 3, and 4, the new outflow is included in *γ*^*′*^, whose index is compared to *γ*, to satisfy the outputcompleteness. In cases 1, 2, and 3, there is a new cycle constructed by {3, 4, outflow-of-B} in the network after the addition of the outflow of B, and it is counted in the substructure of only case 3. In cases 4 and 5, the sum of concentrations of A, B, and C is conserved in time before the addition of the outflow of C, and it is counted in both substructures of cases 4 and 5. See the Methods section C. for the detailed definition of the number of cycles and conserved quantities.

According to the table, when *γ* does not include a conserved quantity constructed from *m*^*^, cases 1, 2, or 3 can occur and the index of *γ* decreases by one or does not change. If *γ* contains chemicals that construct a conserved quantity with *m*^*^, then case 4 or 5 occurs, and the value of index *χ* (*γ*) increases or does not change. There are more than 2^|*M*|^ − 1 output-complete substructures in the network, and this result enables us to determine the change in the index of those substructures caused by the addition of an outflow from the structure of the network.

Cases 1, 2, and 3 can arise when the output-complete substructure *γ* does not contain conserved quantities constructed from *m*^*^ and other chemicals. In case 1, the substructure *γ* contains neither *m*^*^ nor conserved quantities constructed from *m*^*^. Under this condition, no new cycles are created in *γ* because the new outflow is not present in *γ*^*′*^. Therefore, all terms in the index *χ* (*γ*) are not affected by the modification, and *χ* (*γ*) − *χ* (*γ*^*′*^) = 0 holds. For example, the substructure ({*A*}, {2}) in the network shown in Fig. 2C remains a buffering structure before and after the addition of the outflow of B.

Case 2 is induced when *γ* includes *m*^*^, no conserved quantity, including *m*^*^, exists in the substructure, and the addition of the outflow does not create a new cycle in *γ*^*′*^. In this case, since the number of reactions increases by one, while the number of chemicals, cycles, and conserved quantities remain unchanged, the change in the index is 1. For example, in the network shown in Fig. 2C, the subnetwork ({*B* }, {3}) is a buffering structure in the original network, but ({*B*, 3}, {*outflow*-*of* -*B*}) is not a buffering structure in the new network with the outflow of B, and the modification causes a drastic change in the sensitivity.

In case 3, *γ* contains *m*^*^, no conserved quantity, including *m*^*^, exists in the substructure, and the addition of the outflow creates a new cycle within *γ*^*′*^. In this case, the number of reactions and cycles both increase by one, resulting in no change in the index. In case 3 network of Fig. 2C, the substructure ({*B, C* }, { 3, 4}) is a buffering structure in the original network, and ({*B, C* }, {3, 4, *outflow*-*of* -*B* }) is also a buffering structure in the new network with the outflow of B.

Cases 4 and 5 can occur when *γ* contains a conserved quantity constructed from *m*^*^ and other chemicals. In both cases, the conserved quantity constructed from *m*^*^ changes in time after the addition of the outflow of *m*^*^, and the number of conserved quantities decreases by one. Since the number of conserved quantities and cycles do not change simultaneously by the addition of the outflow (for the proof, see the Methods section D.), the number of cycles is not affected in those cases.

Case 4 occurs when *γ* contains *m*^*^, and *m*^*^ constructs a conserved quantity. In this case, the number of reactions increases by one and the number of conserved quantities decreases by one, resulting in no change in the index *χ* (*γ*). In case 4 network in Fig. 2C, the index of the substructure ({*C* }, {3, 4}) is -1 before the addition of the outflow of C, and the index of the substructure ({*C* }, {3, 4, *outflow*-*of* -*C* }) is -1.

Case 5 is induced when *m*^*^ does not exist in *γ*, and other chemicals that construct a conserved quantity with *m*^*^ are included in *γ*. If the condition is satisfied, the number of conserved quantities decreases by one, and the index *χ* (Γ) increases by one. In case 5 network of Fig. 2C, the index of the substructure ({*A*}, {1, 2}) is -1 before the addition of the outflow of C, but after the addition, the index is 0 and the substructure is a buffering structure.

The above classification allows us to predict the change in buffering structures in the focal system caused by the addition of outflows from the original network as follows. First, list all substructures with index 0 or -1 in the original network. Next, for each substructure, calculate which of the five cases follows. The substructures whose indices change from 0 to -1 are buffering structures that are lost by the structural modification, and the substructures whose indices change from -1 to 0 are new buffering structures.

### Analysis of central metabolic system

Using the above result, we examined how structural alterations to the central metabolic system affect cellular metabolism. To create an original system, we integrated KEGG mouse pathways: glycolysis (mmu00010), TCA cycle (mmu00020), and pentose phosphate pathway (mmu00030). We also added inflows and outflows of several metabolites to guarantee the existence of a positive steady-state of the system (Fig. 3A and Table S1). The network contained 48 chemicals, 102 reactions, and 54 cycles, and there were no conserved quantities in the system. In the following, we first explain the sensitivity of this network system based on buffering structures and demonstrate that the effect of network alternation on the sensitivity could be captured by the changes in the indices of output-complete subnetworks, particularly the loss or creation of buffering structures.

**Figure 3:**
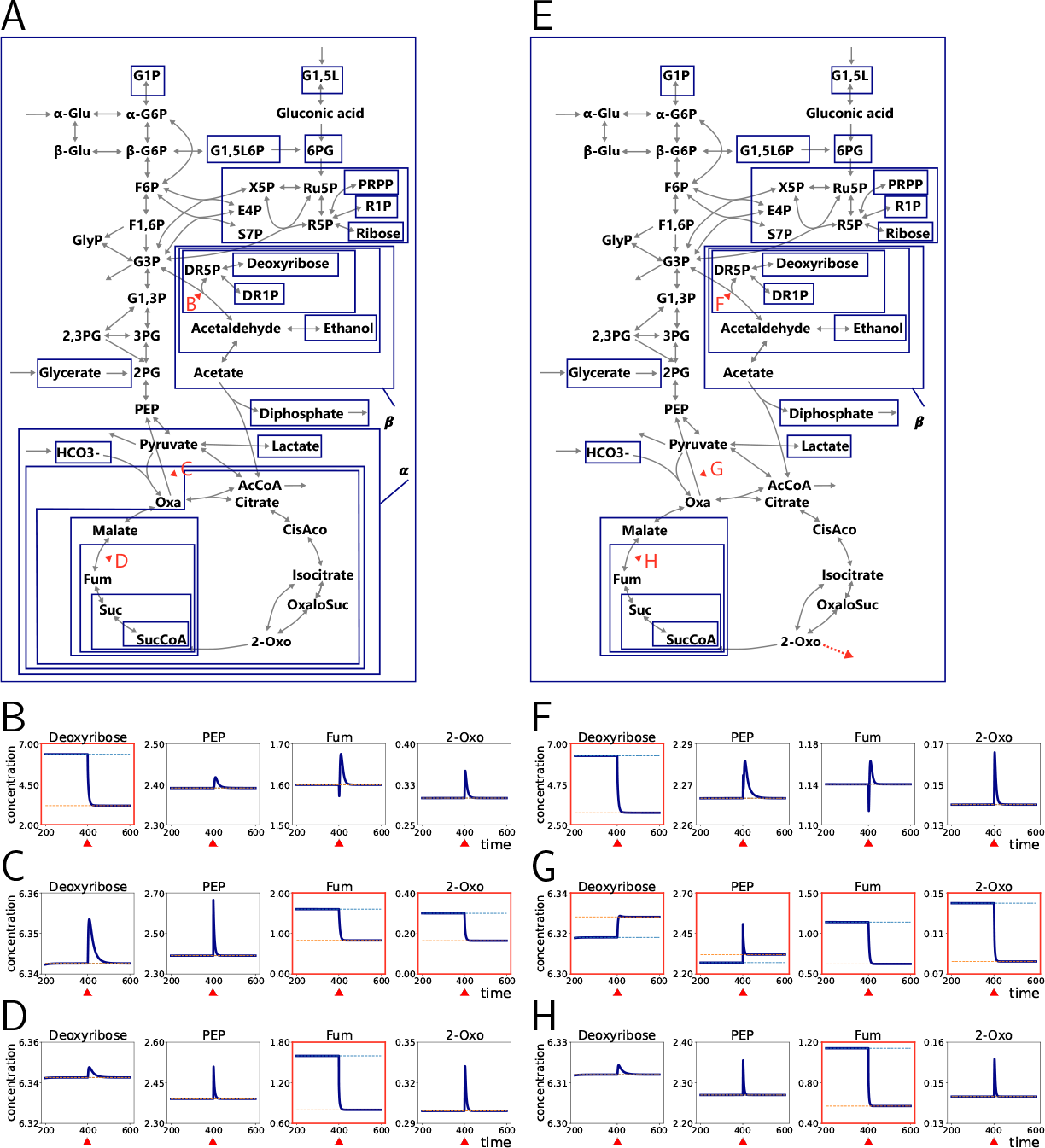
(A) Central metabolic system of mice obtained from KEGG. Red arrowheads indicate the reactions that are perturbed in the numerical calculation. Blue rectangles indicate the buffering structures in the network. (B-D) Numerical simulation results of the dynamics in the network (A) based on mass-action kinetics. The concentrations of four chemicals (Deoxyribose, PEP, Fum, and 2-oxo) are shown. At *t* = 400, where the system is in a steady state, the rate parameter of a reaction indicated by one of the red arrowheads in (A) is varied. The perturbed reactions are (B) G3P→ DR5P + Acetaldehyde, (C) Oxa→ PEP, and (D) Fum → Malate, respectively. The frame of the numerical simulation result is colored red if the steady state value differs before and after the perturbation to the reaction parameter. (E) Buffering structures in the network with the new outflow of 2-oxoglutarate. (F-H) Numerical simulation of the dynamics of the network shown in (E) based on mass-action kinetics. The rate parameter of a reaction indicated by one of the red arrowheads is varied at *t* = 400. The perturbed reactions are (F) G3P → DR5P + Acetaldehyde, (G) Oxa → PEP, and (H)Fum → Malate, respectively. The frame of the numerical simulation result is colored red if the steady state value differs before and after the perturbation to the reaction parameter.

We identified 47 buffering structures within the system by performing the SSA on this system, as shown in Table S2 and by blue rectangles in Fig. 3A. They were found based on the sensitivity obtained by the numerical implementation of the SSA in the Methods section F.. Some of these structures correspond to functional pathways in databases. Specifically, the output-complete substructure *α* and *β* in Fig. 3A correspond to pathways referred to as the TCA cycle and the pentose phosphate pathway, respectively [17]. Because the TCA cycle is a buffering structure, the effect of modulation to reaction parameters within the TCA cycle (e.g., Oxa→ PEP) is expected to be limited to the steady-state chemical concentrations and reaction rates within the TCA cycle. We conducted numerical simulations of dynamics with assumed reaction functions to confirm the expectations from the SSA (Fig. 3B-D, S1). Our simulations showed that modulating the parameter of a reaction in the TCA cycle (Oxa→ PEP) affects only the concentration of chemicals inside the TCA cycle, such as Fumalate and 2-oxoglutarate, and not the concentration of chemicals outside the TCA cycle, such as deoxyribose and PEP, as shown in Fig. 3C, S2. Since the reaction Fum → Malate is contained within a smaller buffering structure inside the TCA cycle, modulating the parameter of this reaction only affects the concentrations of some chemicals within the TCA cycle, including Fumalate, but not the others, such as 2-oxoglutarate (Fig. 3D, S3). The reaction G3P→ GR5P + Acetaldehyde is contained within the buffering structure corresponding to the pentose phosphate pathway. Therefore, activating this reaction only affects the concentration of some chemicals within this buffering structure, including deoxyribose, and does not affect the concentration of chemicals outside the structure (Fig. 3B, S1). This result suggests that in the original system, the concentration of chemicals in the TCA cycle can be regulated by modulation of the expression levels of enzymes catalyzing reactions in the structure without affecting the concentration of chemicals in other pathways, such as the glycolysis pathway and pentose phosphate pathway.

Certain leukemias, gliomas, and other cancer cells have been observed to exhibit a reaction that converts 2-oxoglutarate, a component of the TCA cycle, into a cancer-specific metabolite as a result of gain-of-function mutations [20, 21, 22, 23]. To investigate this phenomenon, we analyzed the changes in buffering structures of the central metabolic system caused by the addition of the outflow of 2oxoglutarate based on which case a buffering structure follows (Fig. 3E, S2). Since 2-oxoglutarate does not contribute to any conserved quantities in the central metabolic system, the addition of the outflow of 2-oxoglutarate leads to changes in the index of each substructure either in case 1, 2, or 3, as shown in Fig. 2 A, B. Therefore, there is no possibility for substructures with an index of -1 to become new buffering structures, and all changes in buffering structures can be captured by simply focusing on the existing ones.

We calculated the index change of buffering structures induced by the addition of the outflow of 2oxoglutarate using Table 2A (Table S2). Out of 47 buffering structures, 38 do not contain 2-oxoglutarate and follow case 1 with no index change. Since eight buffering structures, including the TCA cycle, contain 2-oxoglutarate but no cycle appears in them, case 2 is induced. Case 3 was induced in the one remaining buffering structure because it contains 2-oxoglutarate, and a new cycle is created by the structural modification. Since the TCA cycle is not a buffering structure after the addition of the outflow, modulations to reaction parameters of the reaction in the TCA cycle (Oxa→ PEP) are expected to affect the concentration of chemicals outside the TCA cycle. In contrast, smaller buffering structures within the TCA cycle and the pentose phosphate pathway follow case 1, therefore they remain buffering structures, and the effect of perturbation to inside reactions is also confined in them after the structural modification.

Numerical simulations show that the activation of the reaction in the TCA cycle (Oxa→ PEP) affects the steady-state concentration of chemicals outside the TCA cycle (deoxyribose and PEP) after the addition of the outflow of 2-oxoglutarate (Fig. 3G, S5). Furthermore, the range of influence by the modulation to the reaction parameter of G3P→ GR5P + Acetaldehyde and Fum → Malate does not change before and after the addition of the outflow of 2-oxoglutarate (Fig. 3F, H, S4, S6). Therefore, it can be concluded that the TCA cycle cannot be regulated independently within the central metabolic system when a reaction that converts 2-oxoglutarate into another chemical that is not in the system is added. Conversely, the controllability of the pentose phosphate pathway and small parts of the TCA cycle is not affected by this structural alteration.

We also identified structural modifications that decrease the index of the TCA cycle and make it not a buffering structure by utilizing our result in reverse. In general, the addition of an outflow to a chemical outside the substructure results in an index change of case 1 or 5 in Fig. 2A. Since there is no conserved quantity in the original system, case 4 or 5 do not occur, and the addition of outflow to chemicals outside the TCA cycle results only in case 1. This implies that the TCA cycle remains a buffering structure even when an outflow of chemicals outside the TCA cycle is added.

In contrast, the addition of the outflow to chemicals within the TCA cycle can induce case 2 or 3 in the TCA cycle. We found that outflow of only lactate creates a new cycle in the TCA cycle and induces case 3, while outflows of other chemicals within the TCA cycle decrease the index by -1 in case 2. In other words, large changes in the sensitivity behavior of the TCA cycle are induced when an outflow is added to chemicals within the TCA cycle except for lactate.

The colors of chemicals in Fig. 4A represent the change in the index of the TCA cycle caused by the addition of the outflow of each chemical. The addition of the outflows of green and red chemicals leads to case 1 in the TCA cycle and case 3, respectively. The addition of the outflow of blue chemicals decreases the index of the TCA cycle based on case 2. We did not consider adding outflows of black chemicals because they originally had an outflow.

**Figure 4:**
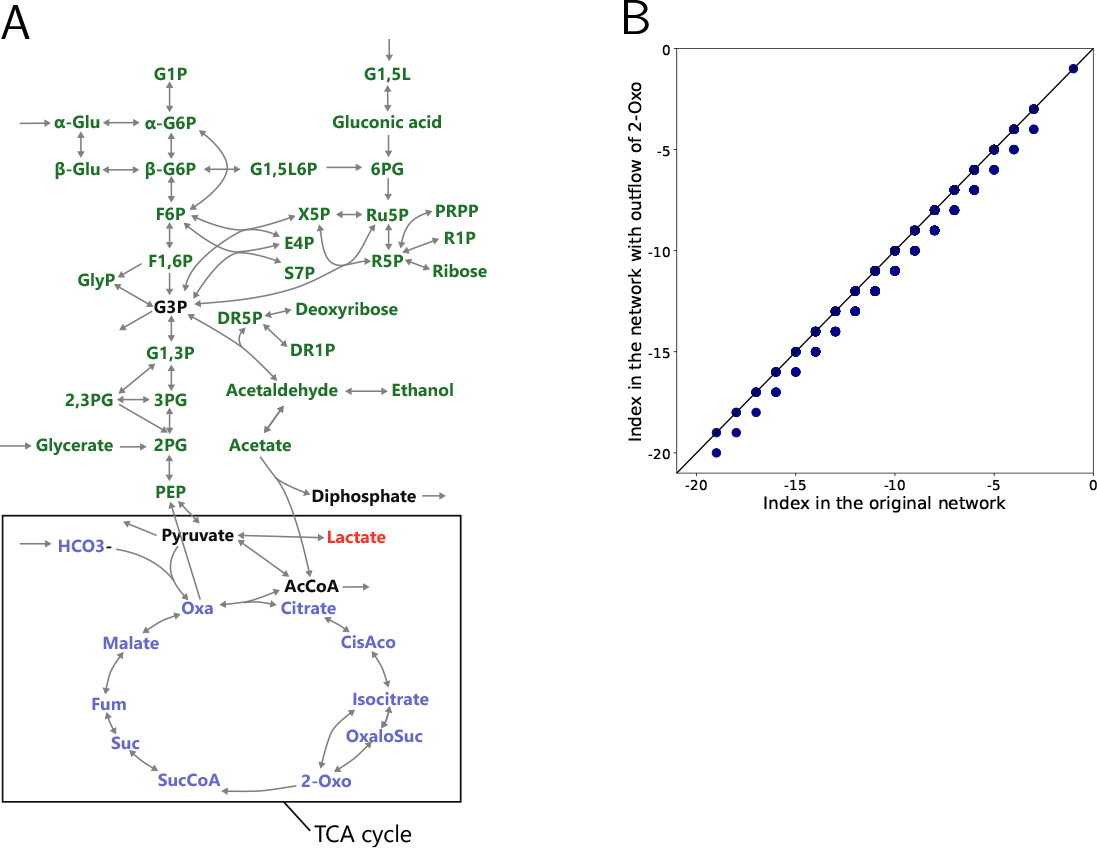
(A) Colors of the nodes indicate how the index of the TCA cycle changes when the outflow is added: green indicates no change in the index in case 1, red indicates no change in case 3, and blue indicates a decrease in case 2. Black nodes are not included in the analysis because they already have an outflow in the original network. (B) Changing of the index of the smallest output-complete substructure containing a randomly selected chemical by adding an outflow of 2-oxoglutarate. The horizontal axis is the index before the addition and the vertical axis is the index after the addition.

To demonstrate that the change in the index of all output-complete substructures caused by the addition of the outflow of 2-oxoglutarate can be identified using the table in Fig. 2A, we calculated the index values of randomly selected output-complete substructures before and after the addition of the outflow of 2-oxoglutarate. First, we randomly selected 10 chemicals from the original system and computed the index of the smallest output-complete substructure that includes the chosen chemicals. Next, we identified the smallest output-complete substructure containing the chosen chemicals on the system with the new outflow and calculated the index values of the substructure on the new network. By repeating the selection of chemicals, we obtained a set of index changes shown in Fig. 4B. Points on the straight line of Fig. 4B indicate cases when the index remains unchanged, while points below the straight line indicate cases when the index decreased by one. Since there are no conserved quantities in the whole system, only cases 1, 2, and 3 can occur, and the change of index is limited to unchanged or decreased.

The numerical simulation of mathematical model based on the KEGG database information does not show any positive steady-state. To solve this issue, we had prepared a network modified from KEGG database by adding some inflows and outflows. To evaluate how this modification affects our result, we investigated buffering structures on the central metabolic network without any modifications, and even in this case, we found that the TCA cycle is a buffering structure. Furthermore, we observed that this buffering structure disappears only when outflow is added to the same metabolites with the main result (metabolites colored in blue in Fig. 4A).

### Analysis of the MAPK signaling pathway

The MAPK signaling pathway is a protein-mediated signaling pathway that is evolutionarily conserved and utilized in many signaling systems. The network shown in Fig. 5A comprises four types of proteins, such as Ras, Raf, Mek, and Erk. Ras has ATP-bounded and ADP-bounded forms, and phosphorylation of Raf, Mek, and Erk is promoted by Ras-GTP, RafP, and MekP, respectively. There are four conserved quantities in this system since the total amount of each protein remains unchanged by ATP binding or phosphorylation.

**Figure 5:**
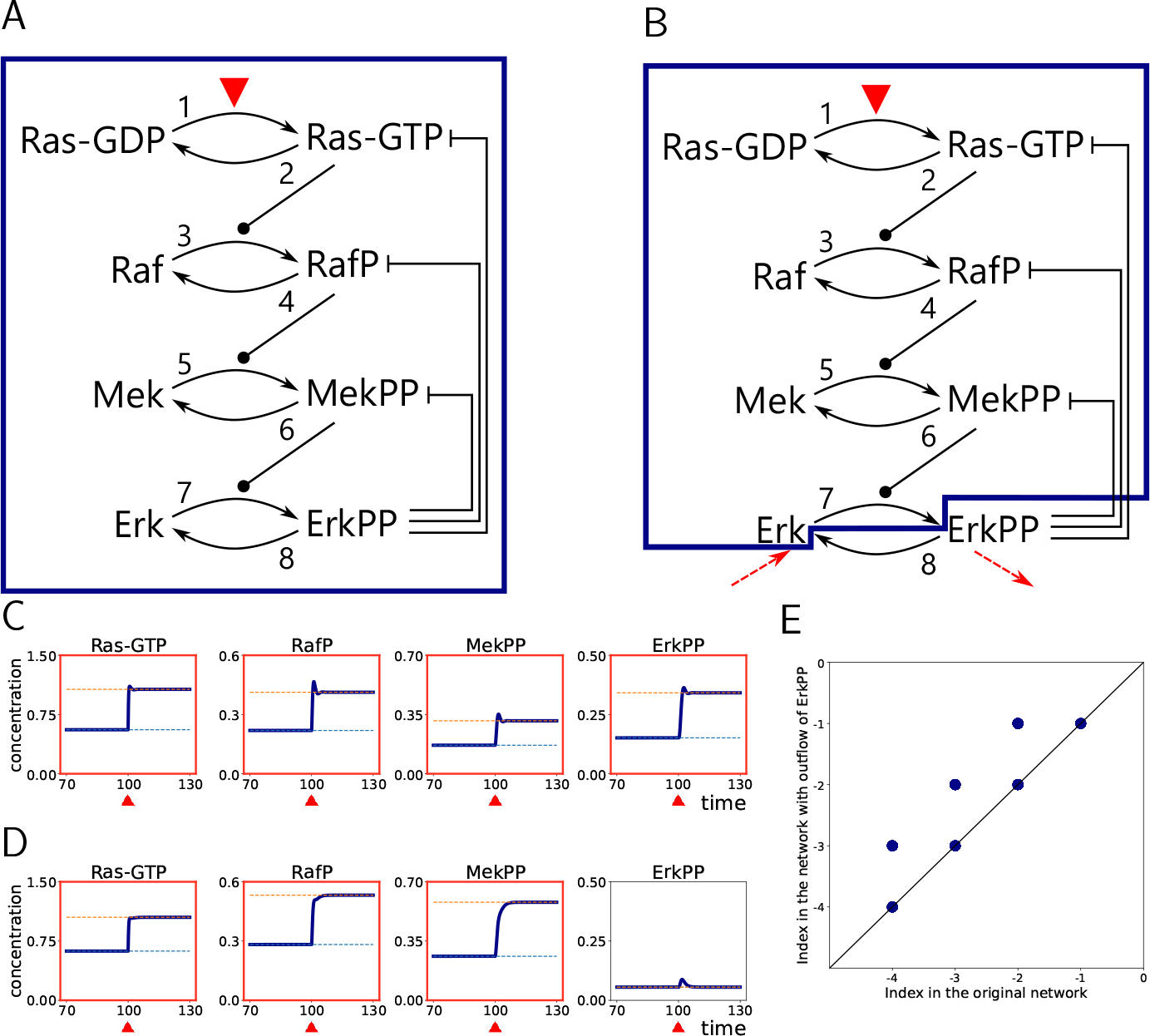
(A) MAPK network. Solid lines represent state transition reactions and dotted lines represent activation or inhibition. The blue square represents the smallest buffering structure containing reaction 1. (B) Network with the outflow of ErkPP. The blue square represents the smallest buffering structure containing reaction 1. (C) Numerical simulation result of the dynamics in the network shown in (A). The parameter of reaction 1 is perturbed at time *t* = 100. At *t* = 100, the dynamics of the network before perturbation reached a steady state. The frame of the numerical simulation result is colored red if the steady state value differs before and after the perturbation to the reaction parameter. (D) Numerical simulation result of the dynamics in the network with the outflow of ErkPP shown in (B). The reaction constants are the same as in (C). The frame of the numerical simulation result is colored red if the steady state value differs before and after the perturbation to the reaction parameter. (E) Changing of the index of the smallest output-complete substructure containing a set of randomly chosen chemicals by the addition of the outflow of ErkPP. Each dot represents an output-complete substructure with randomly chosen chemicals. The horizontal axis is the index before the addition and the vertical axis is the index after the addition.

The blue box in the network in Fig. 5A indicates the smallest buffering structure containing reaction 1, a state transition reaction from Ras-GDP to Ras-GTP. The activation of reaction 1 influences the concentration of all chemicals, including ErkPP, because the buffering structure comprises all chemicals. This is consistent with experimental observations that the MAPK signaling pathway is activated when cells receive signals from outside the cell and regulate the expression of various genes through the nuclear migration of ErkPP. Numerical simulations show that the steady-state concentration of four protein states, including ErkPP, is affected when the parameter of reaction 1 is perturbed (Fig. 5B). Here, although the SSA does not require information on rate functions, and buffering structures are defined only from network structure, specific reaction rate functions are given for numerical simulation, as described in Supplemental Information S4.

Subsequently, we consider the change in the sensitivity of the system by the addition of the outflow of ErkPP, which is the degradation reaction of ErkPP at a rate depending on the ErkPP concentration (Fig. 5C). In the original system, the sum of Erk and ErkPP does not vary over time, but after the addition of the outflow of ErkPP, this quantity is no longer conserved. If an output-complete substructure contains ErkPP, its index does not change due to the structural modification because it follows case 4. When Erk but not ErkPP is present in the substructure, case 5 is induced, and its index increases by one. Substructures that do not contain Erk or ErkPP follow only case 1 since new outflow is not included in them. In summary, the index change of all output-complete substructures is limited to no change or +1; thus, we can capture new or lost buffering structures by focusing on output-complete substructures with an index of 0 or -1.

We examined which case output-complete substructures with the index of 0 or -1 follow (Table S4). In the new system, the substructure shown by the blue rectangle in Fig. 5B is the smallest buffering structure containing reaction 1; thus, the activation of reaction 1 does not influence the concentration of ErkPP. This result indicates that if the degradation rate of ErkPP is high and non-negligible, the activation of reaction 1 does not change the concentration of ErkPP, and signals from outside the cell are not normally transmitted to the nucleus. In other words, the slow degradation of ErkPP is important for the MAPK signaling pathway to function.

Numerical simulations of the dynamics in the new network demonstrate that the concentration of ErkPP temporarily increases when reaction 1 is activated, but its concentration at the steady state remains unchanged before and after the activation of reaction 1 (Fig. 5D). Notably, when the calculation is performed, we added an intake reaction of Erk to the system for the existence of a positive steady state in addition to the outflow of ErkPP, and we confirmed that the addition of this intake reaction to a network with the outflow of ErkPP does not change the index of any output-complete substructures. This example illustrates that the difference in the structure of the network, whether a protein degradation reaction is included or not, can significantly change the sensitivity of the network.

In addition to the buffering structure previously examined, there are more than 2^8^−1 output-complete substructures in the system. We calculated how the index of each output-complete substructure is altered by adding the outflow of ErkPP (Fig. 5E). First, we chose four chemicals randomly from the MAPK network and identified the smallest output-complete substructure containing the selected chemicals. Subsequently, on the new system with an outflow of ErkPP, we found the smallest output-complete substructure that includes the selected chemicals. Then, we calculated the index values of those two substructures. By repeating the choice of chemicals, we obtained a set of index changes shown in Fig. 5E. The points on the straight line indicate cases where the index did not change, the points above indicate cases where the index increased, and the points below indicate cases where the index decreased. As expected from the argument above, there are no points below the line.

## Discussion

We analyzed the effect of network alterations on the sensitivity of a reaction network using the SSA. We found that the addition of an outflow may induce a change in the index of output-complete substructures that exist in a reaction system. The index change is classified into five different patterns depending on the structure of the network of the system. A buffering structure (i.e., a regulatory module) may be created or disappear when the index change is not zero, and it induces a qualitative and macroscopic change in the response of a system to modulations of enzymatic activity.

By focusing on the index change of output-complete substructures, we can predict the creation or loss of buffering structures induced by the addition of an outflow. There is at least one output-complete substructure for all possible chemical subsets, whose total number is 2^|*M*(Γ)|^ − 1. Using our result, we calculated the effect of a structural modification of the network on sensitivity for all those substructures simultaneously. Since the value of the index change is limited to +1 (case 5), -1 (case 2), or no change (case 1, 3, and 4), all new or lost buffering structures can be identified by examining output-complete substructures with the index of +1, -1, or 0 in the original network. We can also identify new outflows that influence the index of selected substructures by utilizing our result in reverse.

We applied our result to biological networks to evaluate the effect of adding a degradation reaction to biological functions. In the central metabolic system of mice, the TCA cycle remains a buffering structure even in the presence of the outflow of chemicals outside the TCA cycle, but its index can decrease when the outflow of chemicals inside the TCA cycle is added to the system. Based on this result, if the degradation rate of those chemicals is not negligible, energy metabolism driven by the TCA cycle cannot be independently regulated from other metabolic pathways. However, it is debatable whether it is biologically plausible that the TCA cycle is a buffering structure.

The numerical calculation in Fig. 3 shows that the perturbation of a reaction from oxaloacetate to PEP (D and H) does not affect the concentration of substances in glycolysis if the TCA cycle is a buffering structure (Fig. 3D), while the perturbation influences glycolysis if the TCA cycle is not a buffering structure (Fig. 3H). The reaction that transforms oxaloacetate to PEP is related to the glucogenesis pathway, which is activated when a cell lacks glucose and needs to synthesize glucose and other substances included in the glycolysis pathway. Considering this, the TCA cycle may not be a buffering structure for the regulation of glucogenesis by the reaction from oxaloacetate to PEP.

In the MAPK network, the addition of an outflow of ErkPP increased the index of a substructure, including Erk and the state transition reaction from Ras-GDP to Ras-GTP, but not ErkPP. The MAPK signal transduction network is activated when cells receive the activation signal of the reaction from Ras-GDP to Ras-GTP, and it leads to the phosphorylation of Erk, resulting in the regulation of gene expressions by ErkPP. From our result, if the degradation rate of ErkPP is high, the extracellular signal cannot change the concentration of ErkPP at a steady state, and the MAPK signaling pathway does not properly function. This indicates that if we could artificially raise the degradation rate of ErkPP, then the extracellular signal would not activate the Erk signaling pathway. Since mutations that constantly activate the Erk signaling pathway are found in some kinds of cancer cells, our result could be used for new cancer therapies [24].

If the index of a buffering structure increases with case 5, and a substructure with a positive index arises, the A-matrix becomes singular, and the entire system cannot be analyzed by the SSA (Method E.). Therefore, we should be careful when adding an outflow of the chemicals that constitute conserved quantities. Although the relationship between the singularity of the A-matrix and the existence of the positive steady state of the system remains elusive, most networks with singular A-matrices do not possess any positive steady states.

Although we focused on the change in sensitivity by the addition of an outflow, the effect of adding a reaction other than the outflow can also be analyzed from the viewpoint of the change in the index of output-complete substructures. For example, consider adding a general reaction in which both substrates and products are present in the original system and check the change in the index of a substructure *γ*. If *γ* does not contain the substrate of the new reaction, the number of reactions and cycles in *γ* remains unchanged, but the number of conserved quantities may change; thus, the index change is *χ* (Γ^*′*^) − *χ* (Γ) = +1 or 0. Therefore, if *γ* is a buffering structure, there is a risk that an output-complete substructure with a positive index will arise, making the entire system unanalyzable.

In this study, we considered the addition of degradation reactions as a modification of a chemical reaction network. In principle, degradation reactions should be present in all chemicals. However, depending on the chemical reaction system and time scale of interest, the degradation reactions of some chemicals may be considerably slow to be interpreted as non-existent. The method we have developed in this paper rather gives criteria for determining whether a degradation reaction should exist for each chemical. For example, in the MAPK signaling pathway, if the degradation reaction of ErkPP exists, the extracellular signal is not transmitted to ErkPP at a steady state. This implies that the degradation reaction of ErkPP should be regarded as negligibly slow compared to the phosphorylation reaction to understand the effect of the growth factor on Erk.

**Table 1:** Summary of notations used in this paper.

|  |  |
| --- | --- |
| $\Gamma, \Gamma'$ | original and altered networks |
| $\gamma, \gamma'$ | output-complete substructures in $\Gamma$ and $\Gamma'$ |
| $\nu$ | stoichiometric matrix of $\Gamma$ |
| $M(\gamma)$ | chemicals in a substructure $\gamma$ |
| $R(\gamma)$ | reactions in $\gamma$ |
| $N_{cyc}(\gamma)$ | number of cycles in $\gamma$ defined in Eq. 4 |
| $N_{cons}(\gamma)$ | number of conserved quantities in $\gamma$ defined in Eq. 5 |
| $\chi(\gamma)$ | index of $\gamma$ defined in Eq. 1 |

Because most biological networks are large and complex, in typical theoretical studies, a subsystem of interest is selected, and a mathematical model is developed for that subsystem. However, it is often not tested on how robust the obtained theoretical conclusions are to the choice of a subsystem. In this study, we established a general rule that determines how the system’s behavior changes with the addition of outflows. This rule provides a systematic understanding of how the selection of subnetworks affects sensitivity. If experimental data on sensitivity is available, by examining the consistency between the experimental data and the theoretical expectations, we can verify whether the subsystem is appropriately chosen or can predict whether unknown outflows should be present or not.

## Methods

### A. The dynamics of chemical reaction system

The time evolution of chemical concentrations *x*_1_, …, *x*_*M*_ in a chemical reaction system is described by an ordinary differential equation system [2, 3, 25, 26], which is expressed as

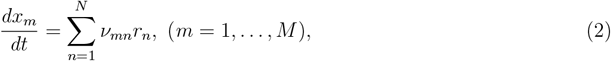

where *r*_1_, …, *r*_*N*_ are the reaction rates and *ν*_*mn*_ is called stoichiometric coefficients. If the stoichiometry of reaction *n* among chemicals *X*_1_, …, *X*_*M*_ is

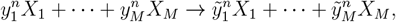

*ν*_*mn*_ is defined as

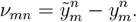

The reaction rate *r*_*n*_ of a reaction *n* depends on the concentration of chemicals ***x*** and a specific parameter *k*_*n*_, and it is described as *r*_*n*_(***x***; *k*_*n*_). The parameter *k*_*n*_ represents the activity or expression level of the enzyme catalyzing the reaction *n*. Mass action law kinetics and Michaelis Menten kinetics are used for modeling reaction rate functions. However, in the SSA, we do not assume any specific kinetics for all reactions. Our formalization includes any regulatory interactions regardless of specific types such as *competitive, noncompetitive*, or *uncompetitive*.

Equation 2 is also described as

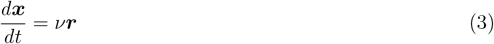

and *ν* is called the stoichiometric matrix.

### B. Mathematical background of the structural sensitivity analysis

The SSA enables us to know how the concentrations of chemicals at a steady state change when reaction parameters are perturbed solely from the network structure of the original system. We consider a steady state of the equation 3. The reaction rate ***r***^*^ at a steady state satisfies *ν****r***^*^ = 0 and is included in ker *ν* = *{****v***|*ν****v*** = 0*}*. When a basis of ker *ν* is selected as *{****c***_1_, …, ***c***_*K*_*}*, ***r***^*^ is expressed using coefficients *μ*_1_, …, *μ*_*K*_,

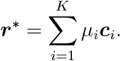

If coker *ν* := *{****v***|***v***^*⊤*^*ν* = 0*}* is nontrivial, the value of ***d***_*l*_ *∈* coker *ν* does not change in time. Thus, a basis of coker 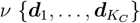 is called conserved quantities.

Let ***A*** be a matrix defined as

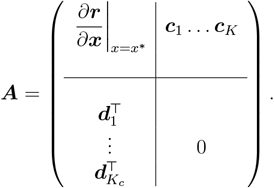

This matrix is always square because the size of *ν* is *M ×N* and rank *ν* = *N* − *K* = *M* − *K*_*c*_ holds from rank-nullity theorem. Notably, whether each component of the matrix is zero or nonzero is determined solely by the network structure. That is,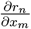 is identically zero when the rate of reaction *n* does not depend on the concentration of chemical *m* and nonzero when it does. The cycles and conserved quantities *c*_*n*_ and *d*_*n*_ are determined from the stoichiometric matrix.

When the A-matrix is regular, the change of concentrations and reaction rates by a parameter modulation is given by

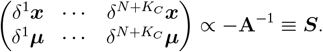

Whether the response of a concentration or reaction rate to a parameter modulation is zero or nonzero can be determined from the distribution of nonzero elements in the S-matrix. The *n*-th column of the S-matrix shows how the steady-state concentration of the chemical and reaction rates change when the parameter of *n*-th reaction *k*_*n*_ is modulated [1]. These are solely determined from the distribution of the zero (non-zero) components of the S-matrix (i.e., the structure of the reaction network).

### C. Mathematical definition of the buffering structure

Let the entire chemical reaction system be Γ, and an output-complete substructure *γ* consisting of a set of chemicals *M* (*γ*) and a set of reactions *R*(*γ*) in Γ. *γ* is output-complete if it contains all reactions whose reactants are in *M* (*γ*). For example, in the network shown in Fig. 2B case 1, a subnetwork ({*A*}, {2} satisfies the output-complete condition.

From equation 3, the stoichiometric matrix *ν* has rows corresponding to chemicals and columns corresponding to reactions. We define the matrix *ν*^*R*(*γ*)^ as a submatrix of *ν* consisting of only the columns corresponding to *R*(*γ*), and *ν*_*M*(Γ)*\M*(*γ*)_ is a submatrix of *ν* constructed from the deletion of rows corresponding to *M* (*γ*). The number of cycles in *γ* is defined as

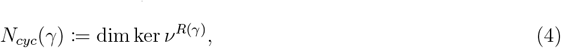

and the number of conserved quantities is expressed as

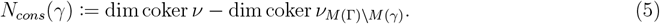

In the case of mono-molecular reaction networks, the cycle in our definition coincides with the cycle in graph theory by introducing a new node that corresponds to the outer part of the system. With such a new node, inflow is expressed as an edge from outside to a chemical, and outflow is an edge from chemical to outside. Note also that *N*_*cons*_(*γ*) is not always equal to dim coker *ν*_*M*(*γ*)_. For example, in the network used at Fig. 2C cases 4 and 5, the sum of concentrations *x*_*A*_ + *x*_*B*_ + *x*_*C*_ remains constant over time. The number of conserved quantities in a substructure (*{A}, {*1*}*) is dim coker *ν* − dim coker *ν*_*B,C*_ = 1, which is different from dim coker *ν*_*A*_ = 0. For a substructure *γ* that is output-complete, we define its index as equation 1. An output-complete substructure *γ* is a buffering structure if *χ*(*γ*) = 0 holds. The modulation of the reaction parameters and conserved quantities in the buffering structure *γ* do not affect the concentration of chemicals outside *γ* [2, 3].

This means that changes in the activity of enzymes catalyzing the reactions in the buffering structure do not alter the concentration of chemicals and rate of reactions outside the buffering structure. Substructures in the network with an index less than 0 do not have the properties of buffering structures. Furthermore, if there is even one substructure with an index greater than 1, the A-matrix reflecting the network structure is singular, making it impossible to analyze sensitivity by the SSA (Methods E.).

### D. Classification of the index change

For a given original system Γ, we consider adding an outflow to a chemical *m*^*^, which does not have outflows originally. The altered system is referred to as Γ^*′*^. For an output-complete substructure *γ* with *M* (*γ*) and *R*(*γ*) in Γ, we define the corresponding output-complete substructure *γ*^*′*^ with *M* (*γ*^*′*^) and *R*(*γ*^*′*^) in Γ^*′*^ as follows: *M* (*γ*^*′*^) in *γ*^*′*^ is identical to *M* (*γ*) in *γ. R*(*γ*^*′*^) is identical to *R*(*γ*) if *M* (*γ*) (or *M* (*γ*^*′*^)) does not contain *m*^*^, while *R*(*γ*^*′*^) is the union of the new outflow and *R*(*γ*) if *M* (*γ*) contains *m*^*^. Obviously, *γ*^*′*^ is output-complete by this definition.

The change in the index is expressed as

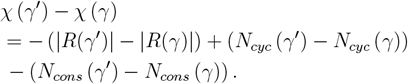

The change in the number of reactions (|*R*(*γ*^*′*^)|− |*R*(*γ*) |) is +1 or 0 depending on whether *R*(*γ*^*′*^) includes the new outflow or not. The new outflow is included in *γ* only when *m*^*^ is in it. The change in the number of cycles (*N*_*cyc*_ (*γ*^*′*^) − *N*_*cyc*_ (*γ*)) is +1 if a new cycle constructed from the new outflow is created in the system or 0 otherwise. The change in the number of conserved quantities *N*_*cons*_ (*γ*^*′*^) − *N*_*cons*_ (*γ*) is -1 or 0 depending on whether substrates that construct conserved quantity with *m*^*^ is included in *M* (*γ*) or not.

1. Cases when *γ* contains *m*^*^ In those cases, the new outflow of *m*^*^ is in the *γ*^*′*^; hence,

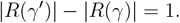

If the new outflow creates a new cycle in the substructure *γ*, the change in the number of cycles (*N*_*cyc*_ (*γ*^*′*^) − *N*_*cyc*_ (*γ*)) is +1, and 0 otherwise. If there is a conserved quantity constructed from *m*^*^ in the original system Γ, the quantity is counted in the number of conserved quantities of the substructure *γ*. The change in the number of conserved quantities (*N*_*cons*_ (*γ*^*′*^) − *N*_*cons*_ (*γ*)) is -1 since the quantity is not conserved after the addition of the outflow. If there is no conserved quantity constructed from *m*^*^ in *γ*, then the change in the number of conserved quantities is 0.
2. Cases when *γ* does not contain *m*^*^

In these cases, the set of reactions of *R*(*γ*^*′*^) is identical to *R*(*γ*); therefore,

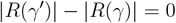

and

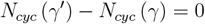

hold. The change in the number of conserved quantities is -1 if there is a chemical that constructs a conserved quantity with *m*^*^, or 0 otherwise.

The above argument for “Cases when *γ* contains *m*^*^” does not exclude a case where the number of reactions increases by one, the number of cycles increases by one, and the number of conserved quantities decreases by one. However, it can be proven that the numbers of cycles and conserved quantities do not change simultaneously.

We show that when the number of cycles increases, the number of conserved quantities does not change. We can use the stoichiometric matrix *ν* of Γ and the unit vector ***e***_*m*_* of which the element corresponding to *m*^*^ is 1 and the others are zero to express the stoichiometric matrix *ν*^*′*^ of Γ^*′*^ as

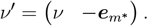

Here, we set the coefficient of *m*^*^ in the chemical equation of the new outflow to 1, but it can take any positive integer without affecting the proof. Since the number of cycles in subsystem *γ* increases, we have

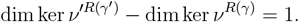

Then from the rank-nullity theorem, we find that

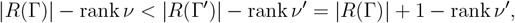

i.e., rank *ν*^*′*^ − rank *ν <* 1. Because rank ≥ *ν*^*′*^ rank *ν*, we conclude that rank *ν* = rank *ν*^*′*^. This means that |*M* (Γ) |− rank *ν* = |*M* (Γ^*′*^)|− rank *ν*^*′*^, that is, the number of conserved quantities in the network remains unchanged. Furthermore, when a vector ***d*** satisfies *ν*^*′⊤*^***d*** = 0, it is also a member of ker *ν*^*⊤*^. Therefore, when we choose a basis of ker *ν*^*′⊤*^, its basis vectors are included in ker *ν*^*⊤*^. Since the rank of ker *ν*^*⊤*^ is equal to that of ker *ν*^*′⊤*^, ker *ν*^*⊤*^ is also spanned by the basis of ker *ν*^*′⊤*^. This leads to

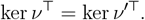

To finalize, we show that the number of conserved quantities in *γ* and that in *γ*^*′*^ is the same. Since ***e***_*m*_* has only nonzero entry in a row corresponding to *m*^*^, *ν*_*M*_*′*_(Γ)\*M*(*γ*)_ has an additional zero column compared to *ν*_*M*(Γ)\*M*(*γ*)_, i.e.,

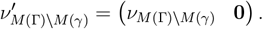

This leads to

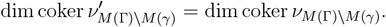

This means that when the number of cycles in a substructure increases by one with a new outflow, the number of conserved quantities of the substructure does not change.

In summary, when *γ* contains *m*\*, either case 2, 3, or 4 occurs, and the case where the number of cycles and the number of conserved quantities simultaneously change cannot occur. When *γ* does not contain *m*^*^, either case 1 or case 5 occurs.

### E. Regularity of A-matrix and substructure with positive index

Suppose that there exists an output-complete substructure *γ*_*s*_ that has a positive index in the original system. Then, we can rearrange rows and columns to transform the A-matrix as

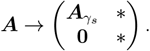

The number of rows of 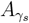 is smaller than the number of columns because the index of *γ*_*s*_ is positive. It means that A-matrix is singular.

### F. Numerical calculation of the S-matrix

The derivatives of reaction rate functions 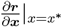appear in the definition of the A-matrix. The SSA does not require the information regarding specific values of 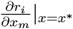and only assumes that 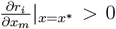 if *m* is the substrate of reaction *i*. For a given reaction network, in principle, qualitative sensitivity (i.e., whether elements of the S-matrix are zero or nonzero (and sometimes whether they are positive or negative)) can be determined by symbolically expressing the inverse of the A-matrix (i.e., the S-matrix) in terms of 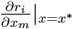.

However, symbolic calculation is computationally expensive for large networks, such as a central metabolic system. Therefore, we assigned real random numbers between 0 and 1 to 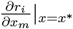 appearing in the A-matrix and then numerically computed the S-matrix to speed up the computation. Therefore, to speed up the computation, we assigned real random numbers between 0 and 1 to 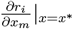appearing in the A-matrix and then numerically computed the S-matrix.

## Data Availability

Details of network structures, list of buffering structures and kinetics for numerical calculations are available in Supplementary Information. The source codes for the computational methods are available on GitHub (https://github.com/hishidagit/NetworkModificationSSA).

## Acknowledgement

A.H. was supported by JSPS KAKENHI (Grant No. 23KJ1324), T.O. was supported by JSPS KAKENHI (Grant No. JP22K03453, JP22K06347) and the RIKEN iTHEMS Program, and A.M. was supported by the CREST program (Grant No. JPMJCR1922) of the Japan Science and Technology Agency (JST) (http://www.jst.go.jp/EN/index.html), Grant-in-Aid for Scientific Research on Innovative Areas (Grant No. 19H05670, 19H03196), and Joint Usage/Research Center program of Institute for Life and Medical Sciences, Kyoto University.

## Author contributions

A.H. performed the data analysis and theoretical proofs. A.M. designed and supervised the research. T.O. developed a new method for the numerical calculation of *S*-matrix explained in Method F.. A.H. wrote the manuscript with the support of A.M. and T.O.

## Notes

### Competing Interest Statement

The authors have declared no competing interest.

