## Supplemental Information for "Patterns of change in regulatory modules of chemical reaction systems induced by network modification"

### Supplementary information for "Determine the impact of network change to the dynamics of chemical reaction systems in cells"

Atsuki Hishida[1], Takashi Okada[2], and Atsushi Mochizuki[1,2,*]

[1]Graduate School of Science, Kyoto University
[2]Institute for Life and Medical Sciences, Kyoto University

November 22, 2023

#### S1 Reactions in the Central Metabolic System

Table S1: Equations of all reactions in central metabolic system of mice shown in Fig. 3A. Metabolic pathways of mice: glycolysis (mmu00010), TCA cycle (mmu00020), and pentose phosphate pathway (mmu00030) were integrated. Inflows were added to G1,5L, Glycerate, Hco3-, DR5P, and $\alpha$-Glu, and outflows were added to Pyruvate, Diphosphate, AcCoA, G3P.

| | substrates | products |
|---|---|---|
| 1 | PEP | Pyruvate |
| 2 | Pyruvate | AcCoA |
| 3 | Acetate | Diphosphate + AcCoA |
| 4 | OxaloSuc | 2-Oxo |
| 5 | 2-Oxo | OxaloSuc |
| 6 | Malate | Oxa |
| 7 | Oxa | Malate |
| 8 | Pyruvate + Hco3- | Oxa |
| 9 | AcCoA + Oxa | Citrate |
| 10 | Citrate | AcCoA + Oxa |
| 11 | Suc | SucCoA |
| 12 | SucCoA | Suc |
| 13 | Oxa | PEP |
| 14 | 2PG | PEP |
| 15 | PEP | 2PG |
| 16 | Lactate | Pyruvate |
| 17 | Pyruvate | Lactate |
| 18 | Isocitrate | 2-Oxo |
| 19 | 2-Oxo | Isocitrate |
| 20 | Acetaldehyde | Acetate |

| | | |
|---|---|---|
| 21 | Acetate | Acetaldehyde |
| 22 | Ethanol | Acetaldehyde |
| 23 | Acetaldehyde | Ethanol |
| 24 | G1P | $\alpha$-G6P |
| 25 | $\alpha$-G6P | G1P |
| 26 | G3P | GlyP |
| 27 | GlyP | G3P |
| 28 | R5P | PRPP |
| 29 | PRPP | R5P |
| 30 | Ribose | R5P |
| 31 | R5P | Ribose |
| 32 | R5P | Ru5P |
| 33 | Ru5P | R5P |
| 34 | R1P | R5P |
| 35 | R5P | R1P |
| 36 | G3P | G1_3P |
| 37 | G1_3P | G3P |
| 38 | DR5P | G3P + Acetaldehyde |
| 39 | G3P + Acetaldehyde | DR5P |
| 40 | F1_6P | GlyP + G3P |
| 41 | GlyP + G3P | F1_6P |
| 42 | Malate | Fum |
| 43 | Fum | Malate |
| 44 | Citrate | CisAco |
| 45 | CisAco | Citrate |
| 46 | 3PG | G1_3P |
| 47 | G1_3P | 3PG |
| 48 | 2_3PG | 3PG |
| 49 | 3PG | 2_3PG |
| 50 | 2PG | 3PG |
| 51 | 3PG | 2PG |
| 52 | G1_5L | Gluconic_acid |
| 53 | 6PG | Ru5P |
| 54 | Ru5P | X5P |
| 55 | X5P | Ru5P |
| 56 | $\beta$-Glu | $\beta$-G6P |
| 57 | $\alpha$-Glu | $\beta$-Glu |
| 58 | $\beta$-Glu | $\alpha$-Glu |
| 59 | S7P + G3P | R5P + X5P |
| 60 | R5P + X5P | S7P + G3P |
| 61 | G1_3P | 2_3PG |
| 62 | 2_3PG | G1_3P |
| 63 | Gluconic_acid | 6PG |
| 64 | $\alpha$-Glu | $\alpha$-G6P |
| 65 | $\alpha$-G6P | $\alpha$-Glu |
| 66 | S7P + G3P | E4P + F6P |

| | | |
|---|---|---|
| 67 | E4P + F6P | S7P + G3P |
| 68 | F6P + G3P | E4P + X5P |
| 69 | E4P + X5P | F6P + G3P |
| 70 | Isocitrate | OxaloSuc |
| 71 | OxaloSuc | Isocitrate |
| 72 | CisAco | Isocitrate |
| 73 | Isocitrate | CisAco |
| 74 | G1_5L6P | 6PG |
| 75 | Suc | Fum |
| 76 | Fum | Suc |
| 77 | $\beta$-G6P | G1_5L6P |
| 78 | $\alpha$-G6P | $\beta$-G6P |
| 79 | $\beta$-G6P | $\alpha$-G6P |
| 80 | $\alpha$-G6P | F6P |
| 81 | F6P | $\alpha$-G6P |
| 82 | DR1P | DR5P |
| 83 | DR5P | DR1P |
| 84 | Deoxyribose | DR5P |
| 85 | DR5P | Deoxyribose |
| 86 | $\beta$-G6P | F6P |
| 87 | F6P | $\beta$-G6P |
| 88 | F6P | F1_6P |
| 89 | F1_6P | F6P |
| 90 | 2-Oxo | SucCoA |
| 91 | Glycerate | 2PG |
| 92 | $\beta$-G6P | $\beta$-Glu |
| 93 | 2_3PG | 2PG |
| 94 | (inflow) | G1_5L |
| 95 | (inflow) | Glycerate |
| 96 | (inflow) | Hco3- |
| 97 | (inflow) | DR5P |
| 98 | (inflow) | $\alpha$-Glu |
| 99 | Pyruvate | (outflow) |
| 100 | Diphosphate | (outflow) |
| 101 | AcCoA | (outflow) |
| 102 | G3P | (outflow) |

Table S2: List of buffering structures in central metabolic system of mice (Fig. 3A). There are subnetworks corresponding to the TCA cycle ($\alpha$, 41), and the Pentose Phosphate Pathway ($\beta$, 36). Note that unions of buffering structures are also buffering structures.

| | substances | reactions | case |
|---|---|---|---|
| 1 | Hco3- | 8 | 1 |
| 2 | SucCoA | 12 | 1 |
| 3 | Lactate | 16 | 1 |
| 4 | Ethanol | 22 | 1 |

| | | | |
|---|---|---|---|
| 5 | G1P | 24 | 1 |
| 6 | PRPP | 29 | 1 |
| 7 | Ribose | 30 | 1 |
| 8 | R1P | 34 | 1 |
| 9 | G1_5L | 52 | 1 |
| 10 | 6PG | 53 | 1 |
| 11 | Gluconic_acid | 63 | 1 |
| 12 | G1_5L6P | 74 | 1 |
| 13 | DR1P | 82 | 1 |
| 14 | Deoxyribose | 84 | 1 |
| 15 | Glycerate | 91 | 1 |
| 16 | Diphosphate | out_Diphosphate | 1 |
| 17 | SucCoA | 11 12 | 1 |
| 18 | Lactate | 16 17 | 1 |
| 19 | Ethanol | 22 23 | 1 |
| 20 | G1P | 24 25 | 1 |
| 21 | PRPP | 28 29 | 1 |
| 22 | Ribose | 30 31 | 1 |
| 23 | R1P | 34 35 | 1 |
| 24 | DR1P | 82 83 | 1 |
| 25 | Deoxyribose | 84 85 | 1 |
| 26 | Suc SucCoA | 11 12 75 | 1 |
| 27 | Suc SucCoA | 11 12 75 76 | 1 |
| 28 | DR1P DR5P Deoxyribose | 38 82 83 84 85 | 1 |
| 29 | Fum Suc SucCoA | 11 12 43 75 76 | 1 |
| 30 | DR1P DR5P Deoxyribose | 38 39 82 83 84 85 | 1 |
| 31 | Fum Suc SucCoA | 11 12 42 43 75 76 | 1 |
| 32 | Fum Malate Suc SucCoA | 11 12 42 43 6 75 76 | 1 |
| 33 | Fum Malate Suc SucCoA | 11 12 42 43 6 7 75 76 | 1 |
| 34 | Acetaldehyde DR1P DR5P Deoxyribose Ethanol | 20 22 23 38 39 82 83 84 85 | 1 |
| 35 | Acetaldehyde DR1P DR5P Deoxyribose Ethanol | 20 21 22 23 38 39 82 83 84 85 | 1 |
| 36 | Acetaldehyde Acetate DR1P DR5P Deoxyribose Ethanol | 20 21 22 23 3 38 39 82 83 84 85 | 1 |
| 37 | E4P PRPP R1P R5P Ribose Ru5P S7P X5P | 28 29 30 31 32 33 34 35 54 55 59 60 66 67 69 | 1 |
| 38 | E4P PRPP R1P R5P Ribose Ru5P S7P X5P | 28 29 30 31 32 33 34 35 54 55 59 60 66 67 68 69 | 1 |
| 39 | 2-Oxo AcCoA CisAco Citrate Fum Isocitrate Malate OxaloSuc Suc SucCoA | 10 11 12 18 19 4 42 43 44 45 5 6 70 71 72 73 75 76 9 90 out_AcCoA | 2 |
| 40 | 2-Oxo AcCoA CisAco Citrate Fum Isocitrate Malate Oxa OxaloSuc Suc SucCoA | 10 11 12 13 18 19 4 42 43 44 45 5 6 7 70 71 72 73 75 76 9 90 out_AcCoA | 2 |

| | | | |
|---|---|---|---|
| 41 | 2-Oxo AcCoA CisAco Citrate Fum Hco3- Isocitrate Lactate Malate OxaloSuc Pyruvate Suc SucCoA | 10 11 12 16 17 18 19 2 4 42 43 44 45 5 6 70 71 72 73 75 76 8 9 90 out_AcCoA out_Pyruvate | 2 |
| 42 | 2-Oxo 2PG 2_3PG 3PG 6PG AcCoA CisAco Citrate DR1P DR5P Deoxyribose E4P F1_6P F6P Fum G1P G1_3P G1_5L6P G3P GlyP Hco3- Isocitrate Lactate Malate OxaloSuc PEP PRPP Pyruvate R1P R5P Ribose Ru5P S7P Suc SucCoA X5P a-G6P a-Glu b-G6P b-Glu | 1 10 11 12 14 15 16 17 18 19 2 24 25 26 27 28 29 30 31 32 33 34 35 36 37 38 39 4 40 41 42 43 44 45 46 47 48 49 5 50 51 53 54 55 56 57 58 59 6 60 61 62 64 65 66 67 68 69 70 71 72 73 74 75 76 77 78 79 8 80 81 82 83 84 85 86 87 88 89 9 90 92 93 out_AcCoA out_G3P out_Pyruvate | 2 |
| 43 | 2-Oxo 2PG 2_3PG 3PG 6PG AcCoA CisAco Citrate DR1P DR5P Deoxyribose E4P F1_6P F6P Fum G1P G1_3P G1_5L6P G3P GlyP Hco3- Isocitrate Lactate Malate OxaloSuc PEP PRPP Pyruvate R1P R5P Ribose Ru5P S7P Suc SucCoA X5P a-G6P a-Glu b-G6P b-Glu | 1 10 11 12 14 15 16 17 18 19 2 24 25 26 27 28 29 30 31 32 33 34 35 36 37 38 39 4 40 41 42 43 44 45 46 47 48 49 5 50 51 53 54 55 56 57 58 59 6 60 61 62 64 65 66 67 68 69 70 71 72 73 74 75 76 77 78 79 8 80 81 82 83 84 85 86 87 88 89 9 90 92 93 in_Glu out_AcCoA out_G3P out_Pyruvate | 2 |
| 44 | 2-Oxo 2PG 2_3PG 3PG 6PG AcCoA CisAco Citrate DR1P DR5P Deoxyribose E4P F1_6P F6P Fum G1P G1_3P G1_5L6P G3P GlyP Glycerate Hco3- Isocitrate Lactate Malate OxaloSuc PEP PRPP Pyruvate R1P R5P Ribose Ru5P S7P Suc SucCoA X5P a-G6P a-Glu b-G6P b-Glu | 1 10 11 12 14 15 16 17 18 19 2 24 25 26 27 28 29 30 31 32 33 34 35 36 37 38 39 4 40 41 42 43 44 45 46 47 48 49 5 50 51 53 54 55 56 57 58 59 6 60 61 62 64 65 66 67 68 69 70 71 72 73 74 75 76 77 78 79 8 80 81 82 83 84 85 86 87 88 89 9 90 91 92 93 in_Glycerate out_AcCoA out_G3P out_Pyruvate | 2 |
| 45 | 2-Oxo 2PG 2_3PG 3PG 6PG AcCoA CisAco Citrate DR1P DR5P Deoxyribose E4P F1_6P F6P Fum G1P G1_3P G1_5L6P G3P GlyP Hco3- Isocitrate Lactate Malate Oxa OxaloSuc PEP PRPP Pyruvate R1P R5P Ribose Ru5P S7P Suc SucCoA X5P a-G6P a-Glu b-G6P b-Glu | 1 10 11 12 13 14 15 16 17 18 19 2 24 25 26 27 28 29 30 31 32 33 34 35 36 37 38 39 4 40 41 42 43 44 45 46 47 48 49 5 50 51 53 54 55 56 57 58 59 6 60 61 62 64 65 66 67 68 69 7 70 71 72 73 74 75 76 77 78 79 8 80 81 82 83 84 85 86 87 88 89 9 90 92 93 in_HCO3- out_AcCoA out_G3P out_Pyruvate | 3 |
| 46 | 2-Oxo 2PG 2_3PG 3PG 6PG AcCoA CisAco Citrate DR1P DR5P Deoxyribose E4P F1_6P F6P Fum G1P G1_3P G1_5L G1_5L6P G3P Gluconic_acid GlyP Hco3- Isocitrate Lactate Malate OxaloSuc PEP PRPP Pyruvate R1P R5P Ribose Ru5P S7P Suc SucCoA X5P a-G6P a-Glu b-G6P b-Glu | 1 10 11 12 14 15 16 17 18 19 2 24 25 26 27 28 29 30 31 32 33 34 35 36 37 38 39 4 40 41 42 43 44 45 46 47 48 49 5 50 51 52 53 54 55 56 57 58 59 6 60 61 62 63 64 65 66 67 68 69 70 71 72 73 74 75 76 77 78 79 8 80 81 82 83 84 85 86 87 88 89 9 90 92 93 in_G1_5L out_AcCoA out_G3P out_Pyruvate | 2 |

| | | | |
|---|---|---|---|
| 47 | 2-Oxo 2PG 2_3PG 3PG 6PG AcCoA Acetaldehyde Acetate CisAco Citrate DR1P DR5P Deoxyribose Diphosphate E4P Ethanol F1_6P F6P Fum G1P G1_3P G1_5L6P G3P GlyP Hco3- Isocitrate Lactate Malate OxaloSuc PEP PRPP Pyruvate R1P R5P Ribose Ru5P S7P Suc SucCoA X5P a-G6P a-Glu b-G6P b-Glu | 1 10 11 12 14 15 16 17 18 19 2 20 21 22 23 24 25 26 27 28 29 3 30 31 32 33 34 35 36 37 38 39 4 40 41 42 43 44 45 46 47 48 49 5 50 51 53 54 55 56 57 58 59 6 60 61 62 64 65 66 67 68 69 70 71 72 73 74 75 76 77 78 79 8 80 81 82 83 84 85 86 87 88 89 9 90 92 93 in_DR5P out_AcCoA out_Diphosphate out_G3P out_Pyruvate | 2 |

#### S2 Results of the numerical simulations of dynamics in the Central Metabolic System

In the numerical simulation of dynamics in the Central Metabolic System, each reaction is assumed to follow mass action kinetics. All reaction parameters are set to 3.0. The time interval of each step in numerical integration was set to $dt = 0.01$ At time point $t = 400$, the parameter of one reaction was increased by 3.0.

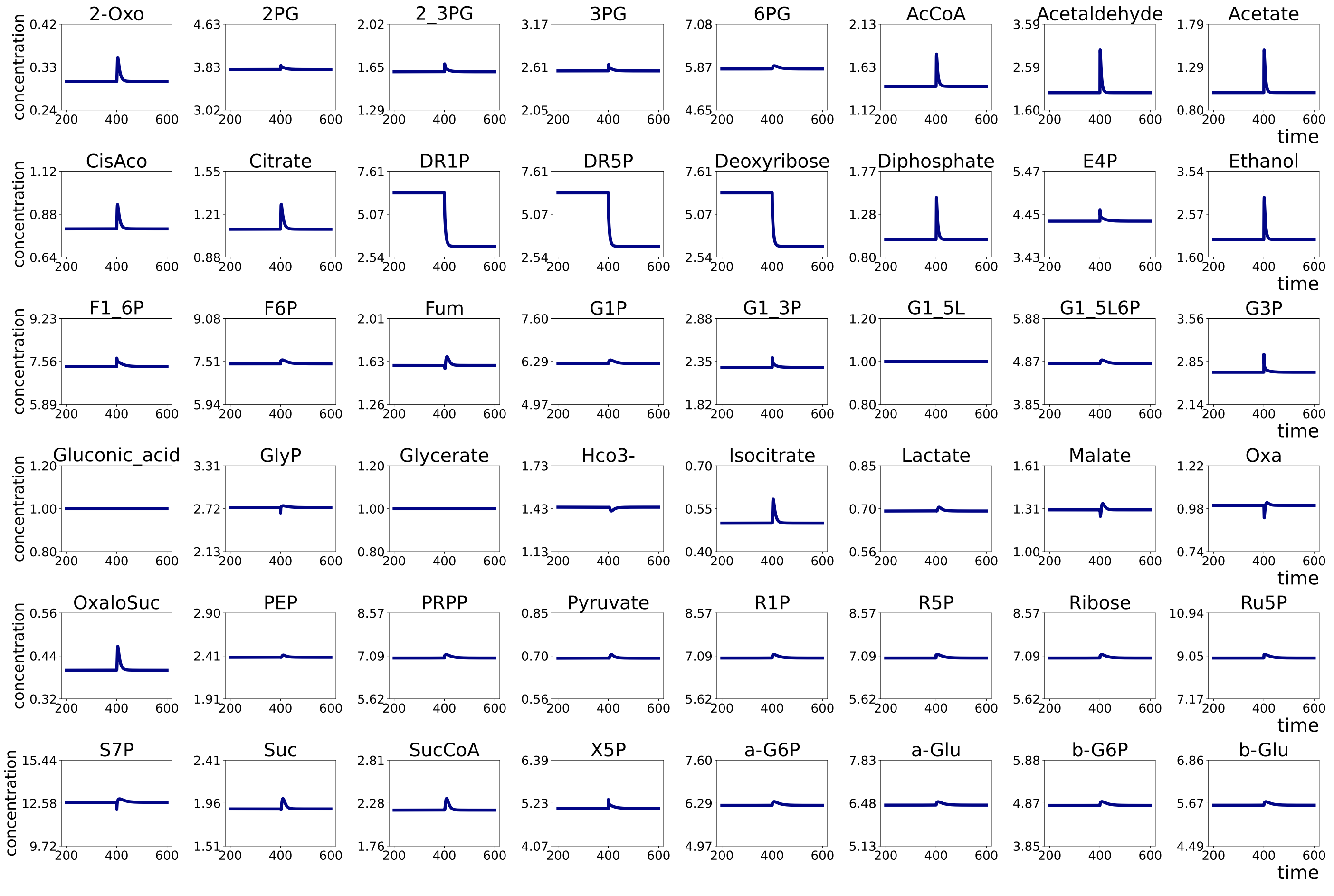


Figure S1: Numerical simulation result of the dynamics of Central Metabolic System. At $t = 400$, the parameter value of reaction 37 : DR5P $\rightarrow$ G3P + Acetaldehyde was increased by 3.0.

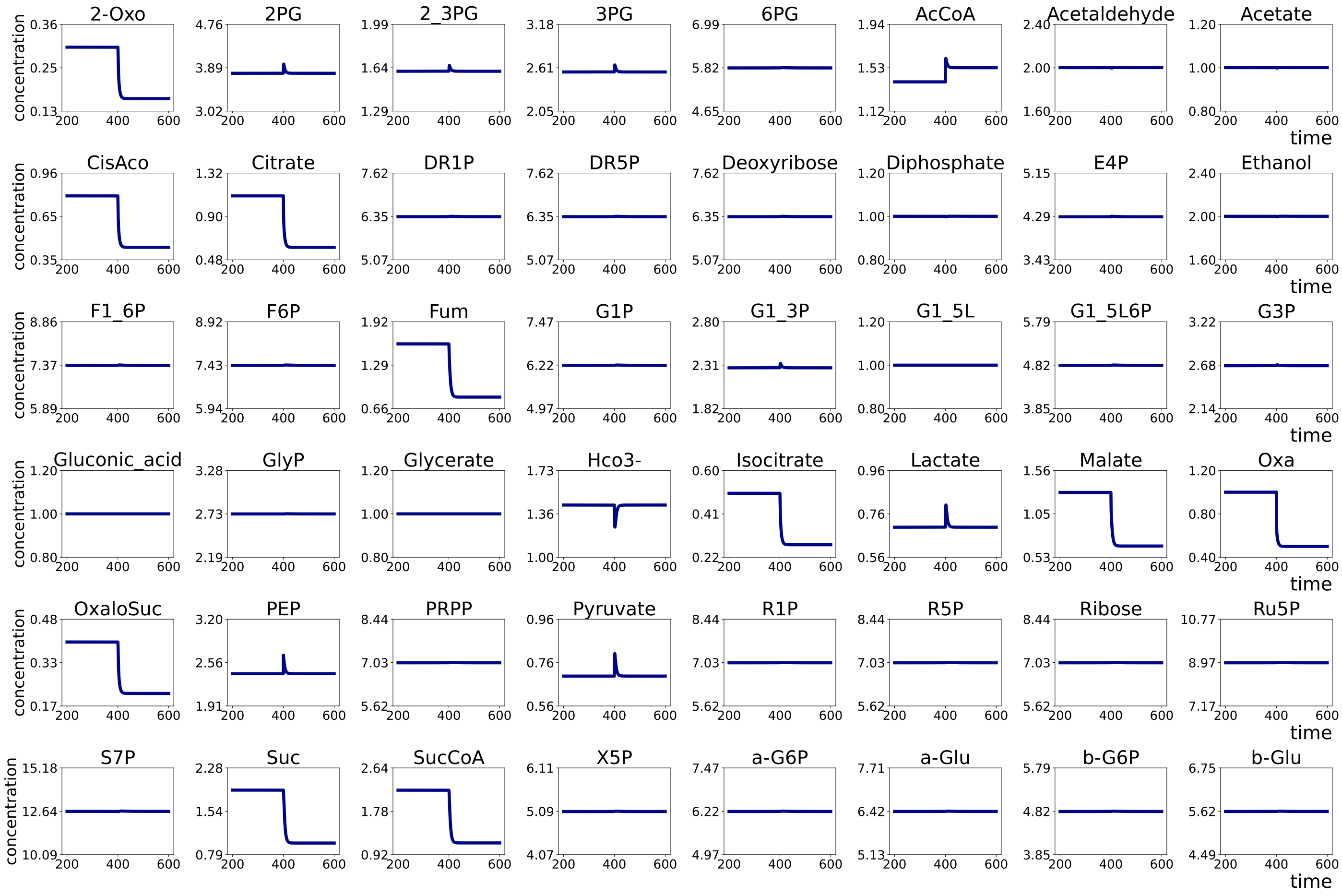


Figure S2: Numerical simulation result of the dynamics of Central Metabolic System. At $t = 400$, the parameter value of reaction 12 : Oxa $\rightarrow$ PEP was increased by 3.0.

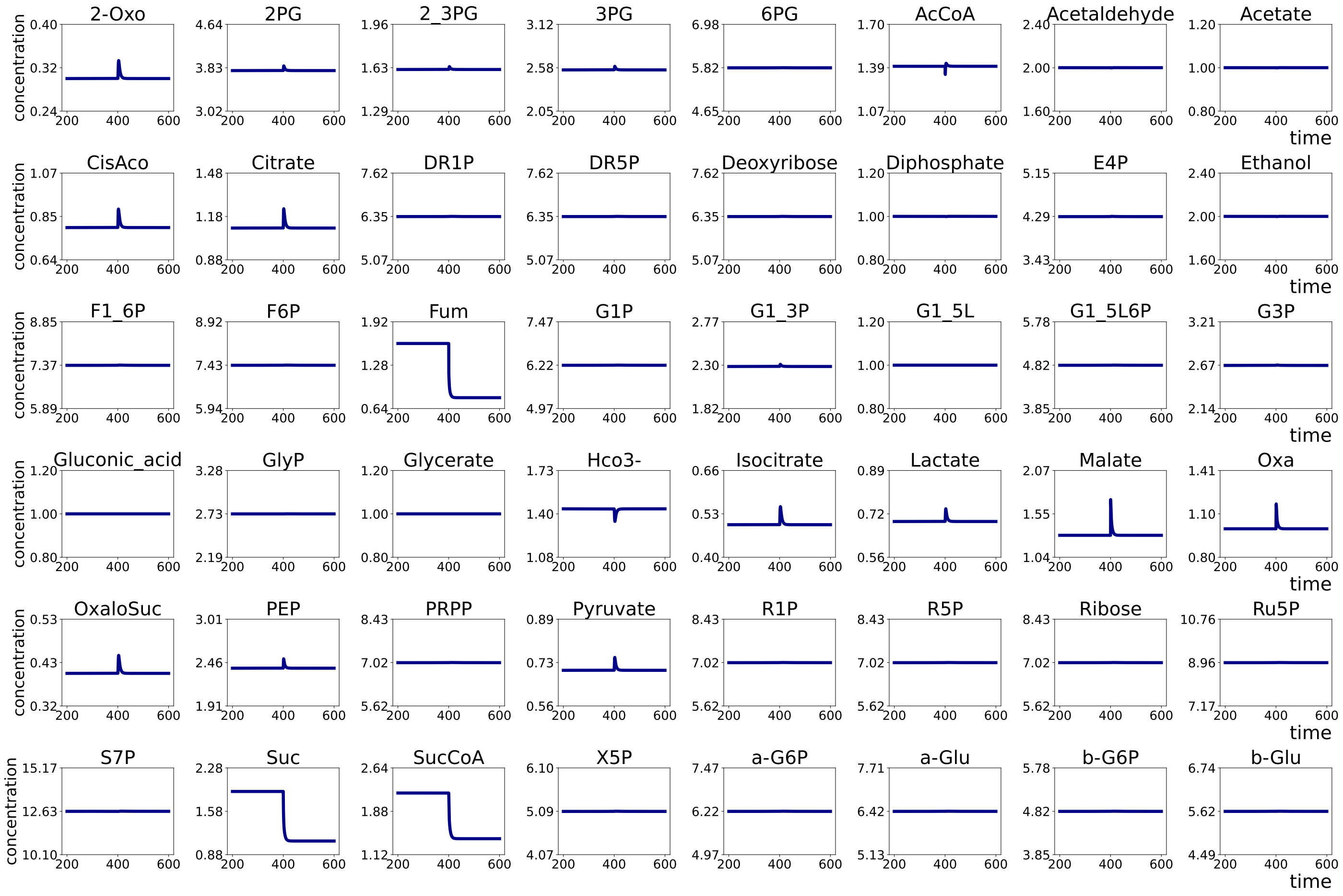


Figure S3: Numerical simulation result of the dynamics of Central Metabolic System. at $t = 400$, the parameter value of reaction 42 : Fum $\rightarrow$ Malate was increased by 3.0.

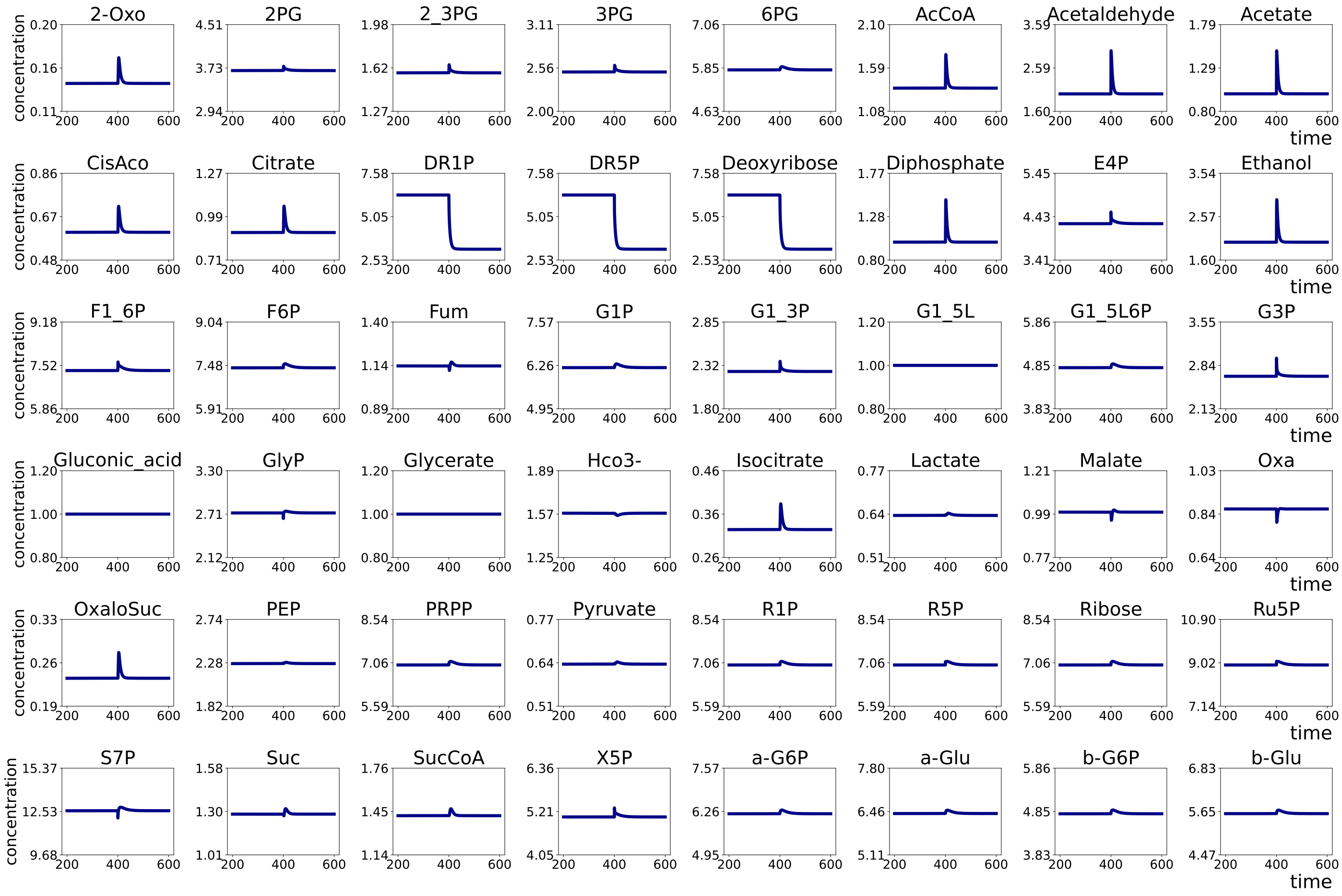


Figure S4: Numerical simulation result of the dynamics of Central Metabolic System with the outflow of 2-Oxoglutarate. At $t = 400$, the parameter value of reaction 37 : DR5P $\rightarrow$ G3P + Acetaldehyde was increased by 3.0.

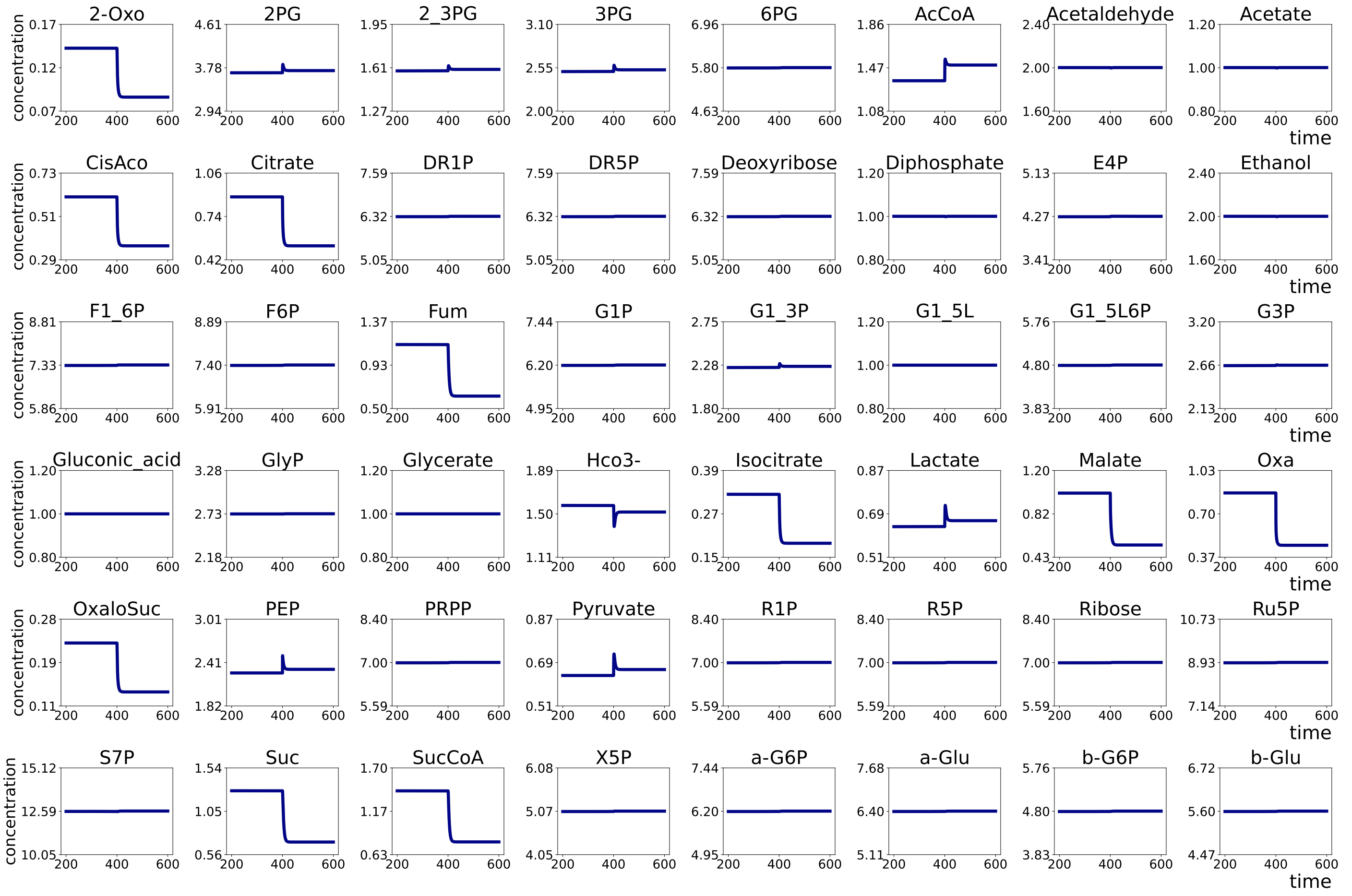


Figure S5: Numerical simulation result of the dynamics of Central Metabolic System with the outflow of 2-Oxoglutarate. At $t = 400$, the parameter value of reaction 12 : Oxa $\rightarrow$ PEP was increased by 3.0.

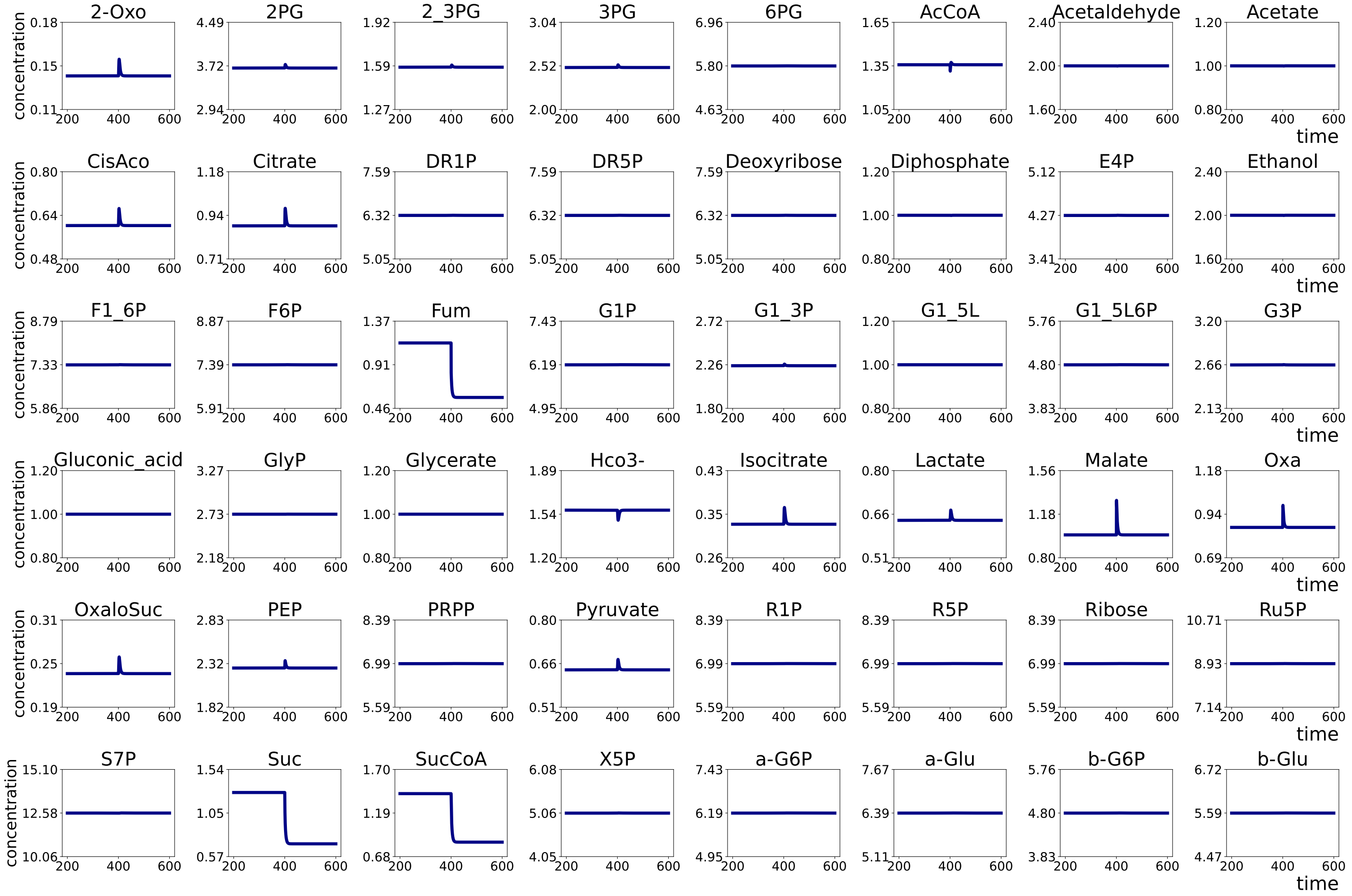


Figure S6: Numerical simulation result of the dynamics of Central Metabolic System with the outflow of 2-Oxoglutarate. At $t = 400$, the parameter value of reaction 42 : Fum $\rightarrow$ Malate was increased by 3.0.

#### S3 Reactions in the MAPK network

Table S3: Reaction equations of all reactions in the MAPK network. Ras-GTP, RafP, and MekPP positively regulates reaction 3, 5, and 7 respectively, and ErkPP negatively regulates reaction 1, 3, and 5.

| | substrates | products |
|---|---|---|
| 1 | Ras-GDP | Ras-GTP |
| 2 | Ras-GTP | Ras-GDP |
| 3 | Raf | RafP |
| 4 | RafP | Raf |
| 5 | Mek | MekPP |
| 6 | MekPP | Mek |
| 7 | Erk | ErkPP |
| 8 | ErkPP | Erk |

#### S4 Results of the numerical simulations of dynamics in the MAPK network

Reaction rate functions of reactions in the MAPK network. Each reaction has a specific parameter $k$, which represents the activity or expression level of the enzyme catalyzing the reaction.

$$f_1(k_1; [\text{RasGDP}], [\text{ErkPP}]) = \frac{k_1 [\text{RasGDP}]}{(K + [\text{RasGDP}])(1 + [\text{ErkPP}])}$$

$$f_2(k_2; [\text{RasGTP}]) = \frac{k_2 [\text{RasGTP}]}{K + [\text{RasGTP}]}$$

$$f_3(k_3; [\text{Raf}], [\text{RasGTP}], [\text{ErkPP}]) = \frac{k_3 [\text{Raf}] [\text{RasGTP}]}{(K + [\text{Raf}])(1 + [\text{ErkPP}])}$$

$$f_4(k_4; [\text{RafP}]) = \frac{k_4 [\text{RafP}]}{K + [\text{RafP}]}$$

$$f_5(k_5; [\text{Mek}], [\text{RafP}], [\text{ErkPP}]) = \frac{k_5 [\text{Mek}] [\text{RafP}]}{(K + [\text{Mek}])(1 + [\text{ErkPP}])}$$

$$f_6(k_6; [\text{MekPP}]) = \frac{k6 [\text{MekPP}]}{K + [\text{MekPP}]}$$

$$f_7(k_7; [\text{Erk}], [\text{MekPP}]) = \frac{k_7 [\text{Erk}] [\text{MekPP}]}{K + [\text{Erk}]}$$

$$f_8(k_8; [\text{ErkPP}]) = \frac{k_8 [\text{ErkPP}]}{K + [\text{ErkPP}]}$$

The default value of each parameter was set to

$$K = 0.5, k_1 = k_2 = k_3 = k_4 = k_5 = k_7 = 3.0, k_6 = k_8 = 1.5$$

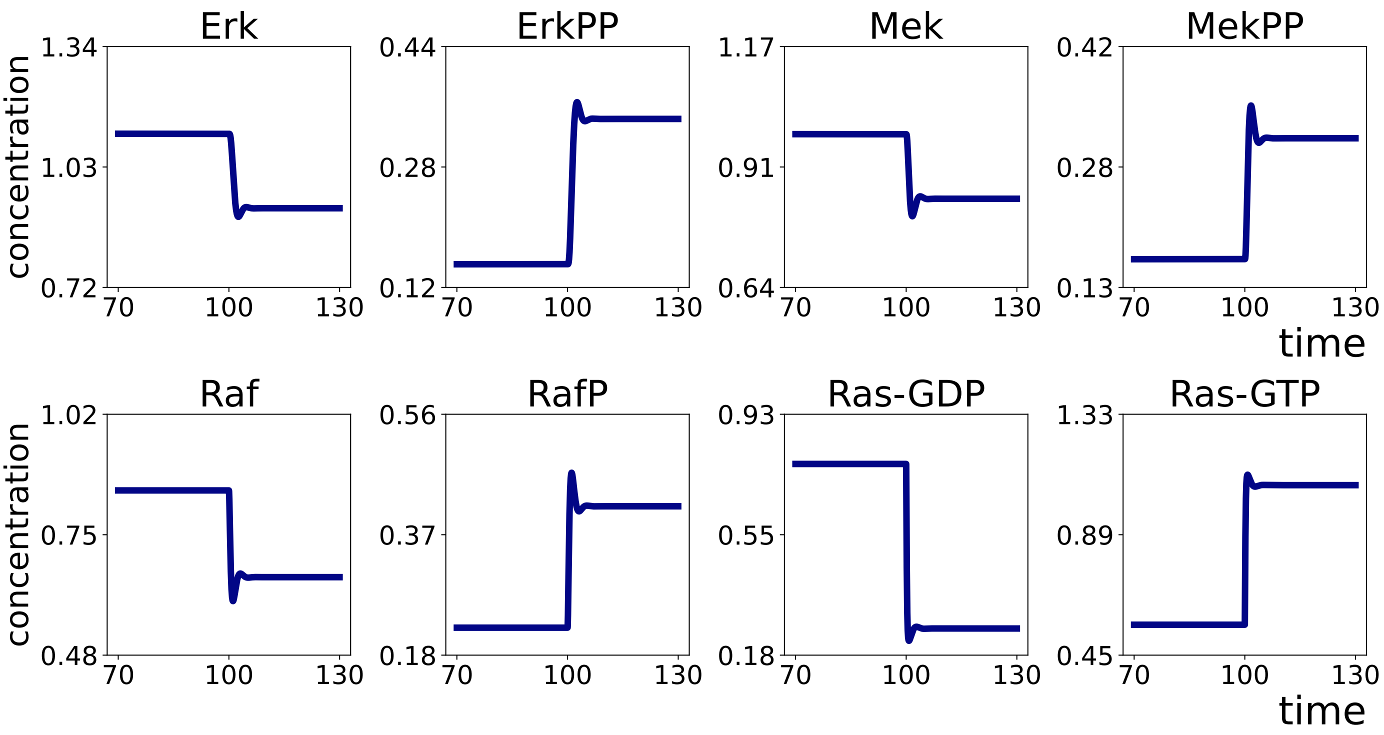


Figure S7: Numerical simulation result of the dynamics of the MAPK signaling pathway. At $t = 100$, the parameter value of the reaction 1: Ras-GDP $\rightarrow$ Ras-GTP was increased by 5.0.

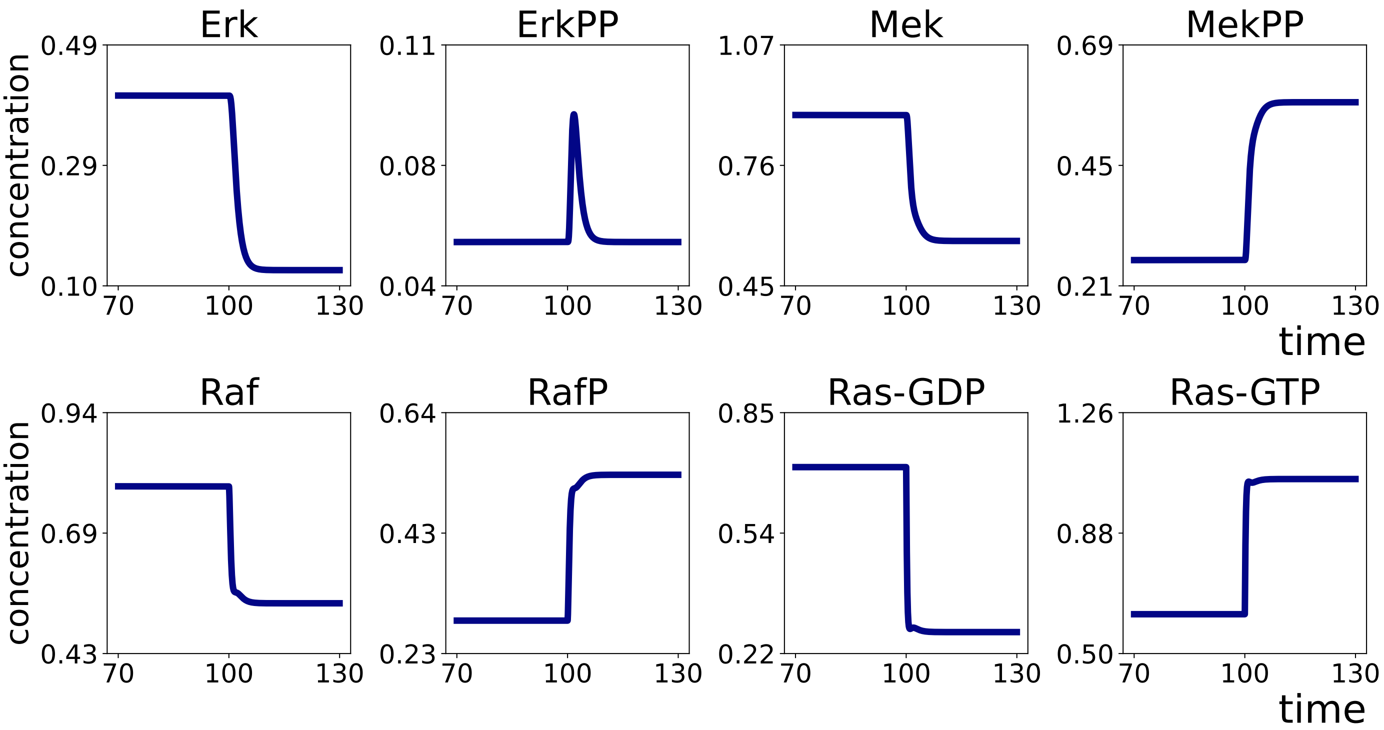


Figure S8: Numerical simulation result of the dynamics of the MAPK signaling pathway with the outflow of ErkPP. At $t = 400$, the parameter value of reaction 1: Ras-GDP $\rightarrow$ Ras-GTP was increased by 5.0.

Table S4: index change of subgraphs with index of -1 or 0 induced by addition of new outflow of ErkPP.

| chemical | reaction | original index | case |
|---|---|---|---|
| Erk,ErkPP,Mek,MekPP,Raf,RafP,Ras-GDP,Ras-GTP | 1,2,3,4,5,6,7,8 | 0 | 4 |
| Erk | 7 | -1 | 5 |
| Mek | 5 | -1 | 1 |
| Raf | 3 | -1 | 1 |
| Ras-GDP | 1 | -1 | 1 |
| Mek,MekPP | 5,6,7 | -1 | 1 |
| Raf,RafP | 3,4,5 | -1 | 1 |
| Ras-GDP,Ras-GTP | 1,2,3 | -1 | 1 |
| Erk,Mek,MekPP | 5,6,7 | -1 | 5 |
| Mek,Raf,RafP | 3,4,5 | -1 | 1 |
| Raf,Ras-GDP,Ras-GTP | 1,2,3 | -1 | 1 |
| Mek,MekPP,Raf,RafP | 3,4,5,6,7 | -1 | 1 |
| Raf,RafP,Ras-GDP,Ras-GTP | 1,2,3,4,5 | -1 | 1 |
| Erk,Mek,MekPP,Raf,RafP | 3,4,5,6,7 | -1 | 5 |
| Mek,Raf,RafP,Ras-GDP,Ras-GTP | 1,2,3,4,5 | -1 | 1 |
| Erk,ErkPP,Mek,MekPP,Raf,RafP | 1,3,4,5,6,7,8 | -1 | 4 |
| Erk,ErkPP,Mek,MekPP,Ras-GDP,Ras-GTP | 1,2,3,5,6,7,8 | -1 | 4 |
| Erk,ErkPP,Raf,RafP,Ras-GDP,Ras-GTP | 1,2,3,4,5,7,8 | -1 | 4 |
| Mek,MekPP,Raf,RafP,Ras-GDP,Ras-GTP | 1,2,3,4,5,6,7 | -1 | 1 |
| Erk,ErkPP,Mek,MekPP,Raf,RafP,Ras-GDP | 1,3,4,5,6,7,8 | -1 | 4 |
| Erk,ErkPP,Mek,MekPP,Raf,RafP,Ras-GTP | 1,2,3,4,5,6,7,8 | -1 | 4 |
| Erk,ErkPP,Mek,MekPP,Raf,Ras-GDP,Ras-GTP | 1,2,3,5,6,7,8 | -1 | 4 |
| Erk,ErkPP,Mek,MekPP,RafP,Ras-GDP,Ras-GTP | 1,2,3,4,5,6,7,8 | -1 | 4 |
| Erk,ErkPP,Mek,Raf,RafP,Ras-GDP,Ras-GTP | 1,2,3,4,5,7,8 | -1 | 4 |
| Erk,ErkPP,MekPP,Raf,RafP,Ras-GDP,Ras-GTP | 1,2,3,4,5,6,7,8 | -1 | 4 |
| Erk,Mek,MekPP,Raf,RafP,Ras-GDP,Ras-GTP | 1,2,3,4,5,6,7 | -1 | 5 |
| ErkPP,Mek,MekPP,Raf,RafP,Ras-GDP,Ras-GTP | 1,2,3,4,5,6,7,8 | -1 | 4 |
